# Exome-wide age-of-onset analysis reveals exonic variants in *ERN1, TACR3* and *SPPL2C* associated with Alzheimer’s disease

**DOI:** 10.1101/2020.01.28.923789

**Authors:** Liang He, Yury Loika, Yongjin Park, Genotype Tissue Expression (GTEx) consortium, David A. Bennett, Manolis Kellis, Alexander M. Kulminski, for the Alzheimer’s Disease Neuroimaging Initiative

## Abstract

Despite recent discovery in GWAS of genomic variants associated with Alzheimer’s disease (AD), its underlying biological mechanisms are still elusive. Discovery of novel AD-associated genetic variants, particularly in coding regions and from APOE ε4 non-carriers, is critical for understanding the pathology of AD. In this study, we carried out an exome-wide association analysis of age-of-onset of AD with ~20,000 subjects and placed more emphasis on APOE ε4 non-carriers. Using Cox mixed-effects models, we find that age-of-onset shows a stronger genetic signal than AD case-control status, capturing many known variants with stronger significance, and also revealing new variants. We identified two novel rare variants, rs56201815, a synonymous variant in ERN1, from the analysis of APOE ε4 non-carriers, and a missense variant rs144292455 in TACR3. In addition, we detected rs12373123, a common missense variant in SPPL2C in the MAPT region in APOE ε4 non-carriers. In an attempt to unravel their regulatory and biological functions, we found that the minor allele of rs56201815 was associated with lower average FDG uptake across five brain regions in ADNI. Our eQTL analyses based on 6198 gene expression samples from ROSMAP and GTEx revealed that the minor allele of rs56201815 was associated with elevated expression of ERN1, a key gene triggering unfolded protein response (UPR), in multiple brain regions, including posterior cingulate cortex and nucleus accumbens. Our cell-type-specific eQTL analysis of based on ~80,000 single nuclei in the prefrontal cortex revealed that the protective minor allele of rs12373123 significantly increased expression of GRN in microglia, and was associated with MAPT expression in astrocytes. These findings provide novel evidence supporting the hypothesis of the potential involvement of the UPR to ER stress in the pathological pathway of AD, and also give more insights into underlying regulatory mechanisms behind the pleiotropic effects of rs12373123 in multiple degenerative diseases including AD and Parkinson’s disease.

## Introduction

Late-onset sporadic Alzheimer’s disease (AD) is a progressive neurodegenerative disorder accounting for 50–70% of all dementia cases in the elderly population [1]. Amyloid β-peptide (Aβ) was the primary component found in the neuritic plaques of AD patient brain, and multiple mutations in the *APP* gene and its related genes (*PSEN1* and *PSEN2*) promoting Aβ production have been identified in familial (early-onset) AD [2–6]. These observations support a causal role of Aβ deposition in the etiology of AD. Familial AD is, however, much rarer than sporadic AD, which is highly prevalent after age 65. Recent genome-wide association studies (GWAS) have identified a large number of genetic variants associated with the risk of late-onset AD [7–13], most of which are located in genes exclusively expressed in microglia (e.g., *TREM2*). These insights suggest the involvement of microglia in the pathology of AD.

Despite recent progress in understanding the biological mechanisms underlying AD, the cellular and molecular activities and causation in the late-onset AD of most common variants discovered in GWAS, including those in *APOE,* remain unclear. Functional links between most of these AD-related loci and genes are still to be determined, although some microglia-related single nucleotide polymorphisms (SNPs) in, e.g., *CD33*, and the *MS4A* gene cluster, are shown to be mediated through *TREM2* [14, 15]. The functional mechanisms of *TREM2* in Aβ uptake by microglia are also complicated, and contradictory biological consequences are observed in mouse models (see, e.g., [16] for a review on this topic). Moreover, adding up the *APOE* variant and other nine identified top SNPs accounts for a small portion (5%) of variation of age-of-onset [17], suggesting that missing genetic mechanisms contribute to this complex disease. We expect that the discovery of additional AD-associated genetic variants will provide more insights into the understanding of the AD pathology.

In this study, we performed an exome-wide association analysis of age-of-onset of AD, in which most genetic variants are rare or low frequency, using an Alzheimer’s Disease Sequencing Project (ADSP) sample of 10,216 subjects in the discovery phase. Rare coding variants often show larger effect size, and their biological consequences are more explicable, but its association analysis is complicated by insufficient statistical power. Although the exome-wide association of AD has recently been explored using AD status [18–20], our rationale is that more AD-related rare variants can be identified using analysis of age-of-onset of AD with a Cox model given emerging evidence from a previous study showing its potential advantage in terms of statistical power [21]. We attempted to replicate significant findings in four other studies, with a meta-analysis sample size of about 20,000 subjects. To understand the biological consequences of the identified SNPs, we explored their influence on regulatory activities and gene expression at tissue and single-cell levels.

We further performed a separate exome-wide association analysis of the age-of-onset of AD by excluding the *APOE ε4* carriers. The overarching goal is to identify novel variants contributing to AD independently of the *APOE ε*4 allele, the strongest single genetic risk factor for AD. Despite quarter Century research on the function of *APOE* gene [22], the primary biological role of this gene in AD pathogenesis remains elusive as the gene and its protein are probably involved in many pathways related to Aβ deposition, Aβ clearance, tau pathology, and neuroinflammation [23]. Our analysis is designed to provide more insights into the AD-related *APOE* biology.

## Results

### Description of the study sample in the discovery phase

In the discovery phase, we carried out an exome-wide association analysis of the age-of-onset of AD using a whole-exome sequencing (WES) sample from the ADSP [24]. We included 10,216 non-Hispanic white subjects (54.86% cases, 58.03% women) after filtering subjects with missing information about sex, AD status, or age-of-onset. The average age-of-onset of AD was 75.4 years (Table S1). We interrogated 108,509 biallelic SNPs with a missing rate < 2% across the subjects and a minor allele count (MAC) > 10. To identify genetic variants associated with the hazards of AD, we conducted three separate analyses. In the first and second analyses, we included all subjects and performed ε*4* allele (coded by the minor allele of rs429358) unconditional (first) and conditional (second) analyses as *APOE ε4* is a well-known strong predictor of AD. That is, we tested two models, differing as to whether the copy of the *APOE ε4* SNP rs429358 was included as a covariate. In the third analysis, we only included 7185 *APOE ε4* non-carriers. Despite this reduction of the sample size, we expect better statistical power by using the age-of-onset analysis. In all analyses, we included sex and a top principal component (PC) as covariates (we examined the top five PCs, and found that only PC2 was associated with AD). As the ADSP sample contains family members, all age-of-onset analyses were performed using Cox mixed effects models implemented in the coxmeg R package [21]. We built a genetic relatedness matrix (GRM) using the ADSP WES data to correct for the relatedness of the subjects. We found that the genomic inflation was controlled in all three analyses (λ=1.04, 1.09, and 1.06) (Fig. S1), comparable to those in [18] using logistic regression models (λ=1.006~1.087).

### Exome-wide analysis of age-of-onset of AD in the discovery phase

In the first analysis (using all subjects without the adjustment for *APOE ε4*), we detected four independent signals passing the exome-wide threshold (p=5e-07) (Fig. 1A, Table S2, Model 1). The most significant SNP was the *APOE ε4*-coding variant rs429358, having a hazard ratio (HR) of 4.37 (p=3.729e-508). The p-value is much more significant than that reported in the largest meta-analysis so far based on AD status (p=5.79e-276) [10]. This result confirms previous findings [25–27] that *APOE ε4* is not only associated with AD status but also substantially decreases its age at onset (Fig. 2A). The three signals outside the *APOE* region are rs75932628 (the R47H mutation) in *TREM2* (HR=2.72, p=2.04e-16), rs7982 in *CLU* (HR=0.887, p=5.98e-08), and rs2405442 in *PILRA* (HR=0.88, p=7.67e-08) (Fig. 1A, Table S2, Model 1). The beneficial association of the missense variant rs7982 in *CLU* was not reported in the previous study of AD status using the same ADSP sample [18]. We observed that the minor allele carriers of rs7982 had lower hazards consistently across a wide age interval (Fig. 2B).

**Figure 1.**
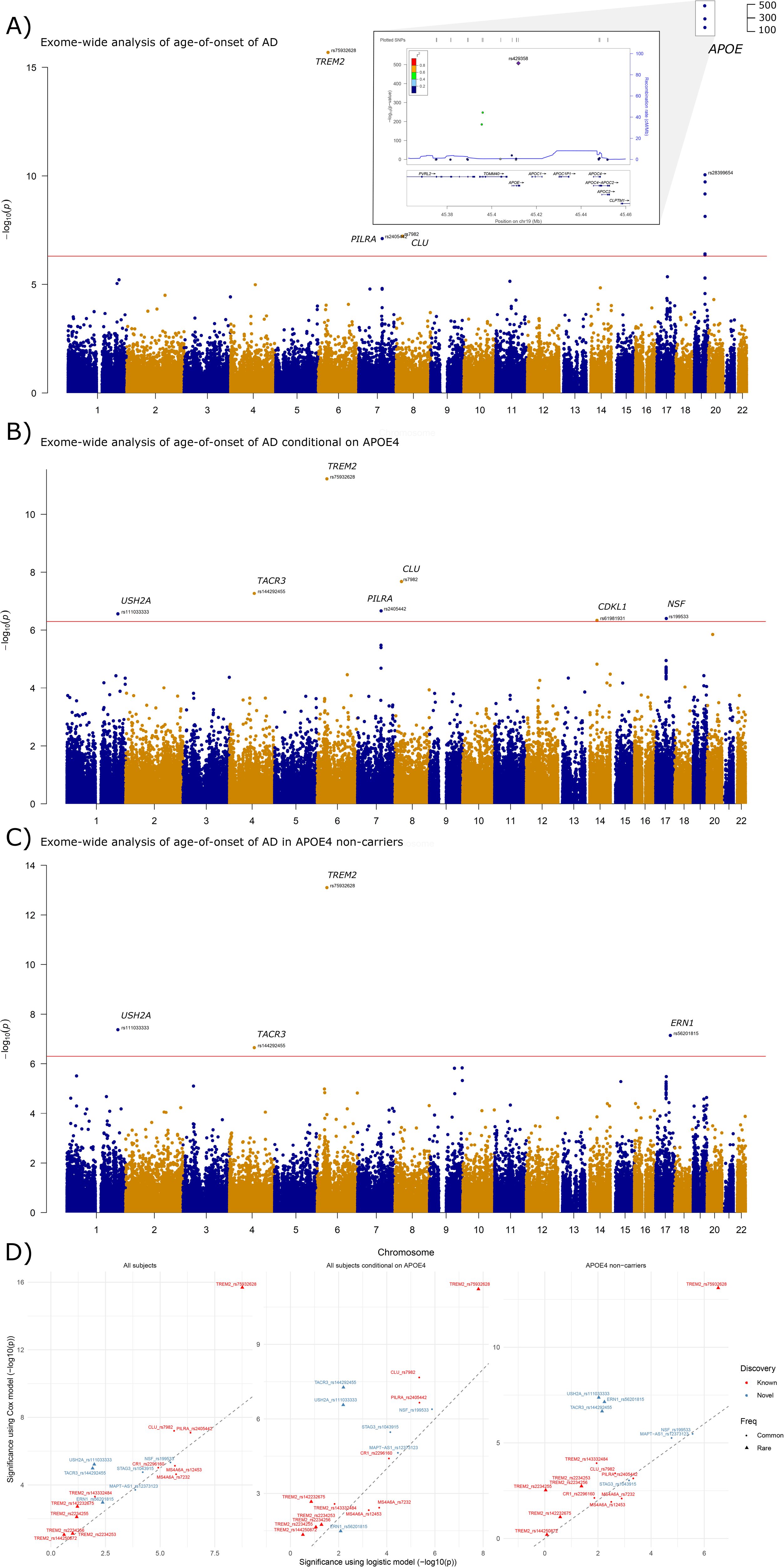
Results of exome-wide association analyses of age-of-onset of AD in the ADSP sample using (A) Model 1: a model with all subjects adjusted for a PC and sex; (B) Model 2: a model with all subjects adjusted for copy of *APOE ε4*, PC2 and sex; (C) Model 3: a model with *APOE ε4* non-carriers adjusted for PC2 and sex. Three top SNPs identified in the *APOE* region using Model 1 were highlighted in the regional plot due to their extremely significant p-values. The red horizontal line is a threshold based on the Bonferroni correction (0.05/100,000=5e-07). (D) Comparison of p-values between a Cox model and a logistic model for well-known AD-related SNPs and newly identified SNPs in this study in Model 1 (left), Model 2 (middle), and Model 3 (right). The same ADSP data and covariates were used to fit the Cox and logistic models.

**Figure 2.**
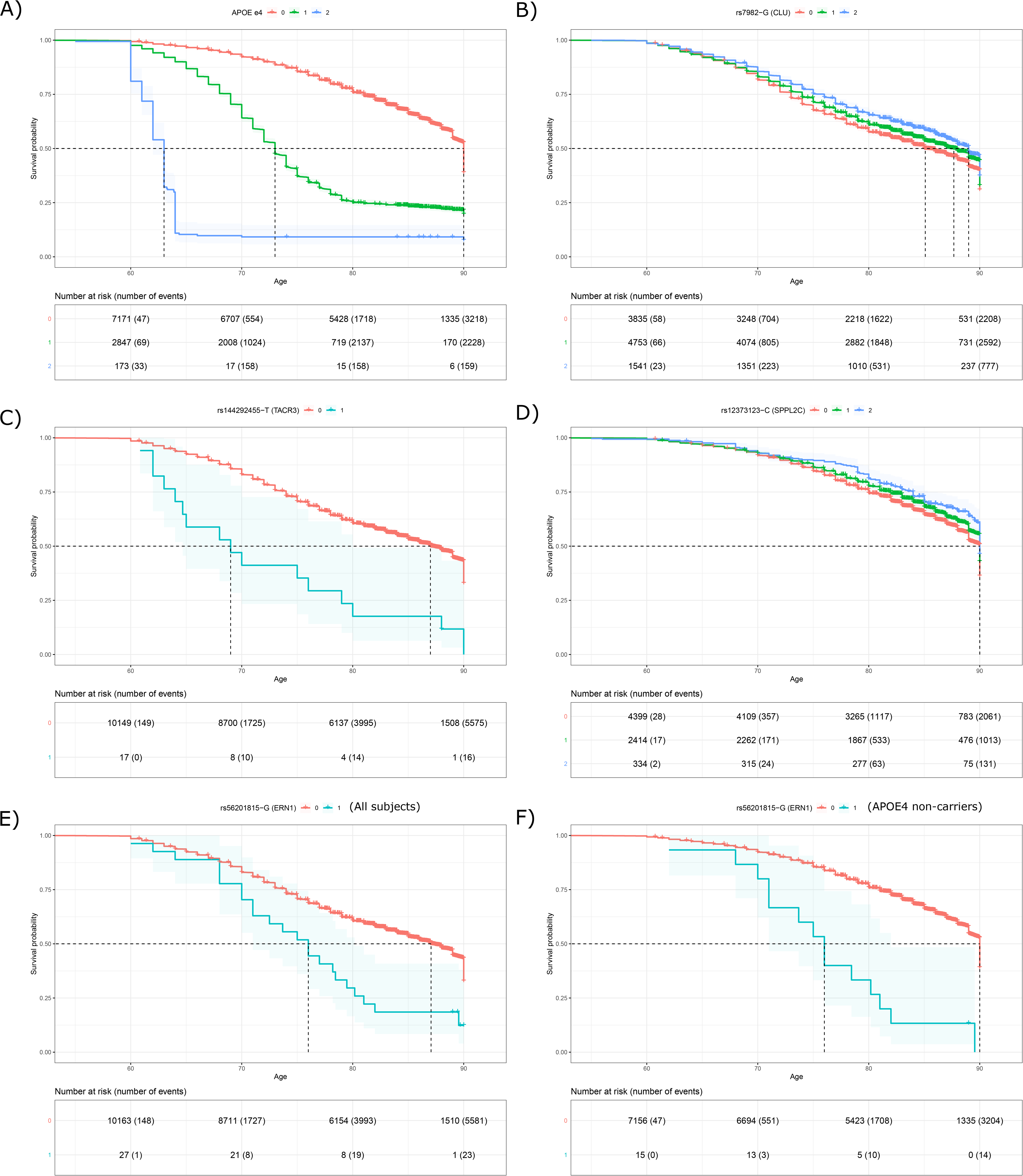
Probability of remaining free of AD (Survival probability) and risk tables in the ADSP sample for genotype groups of (A) *APOE ε4;* (B) rs7982; (C) rs144292455; (D) rs12373123; (E) rs56201815 in all subjects; (F) rs56201815 in *APOE ε4* non-carriers.

Although the R47H mutation in *TREM2* and rs2405442 in *PILRA* were identified in the previous analysis [18], our analysis increased significance for the R47H mutation (p=2.04e-16 vs. 4.8e-12). In addition, we observed well-known AD-associated SNPs among the top hits, including rs12453 in *MS4A6A* (p=7.08e-06), rs2296160 in *CR1* (p=9.05e-06), and rs592297 in *PICALM* (p=5.25e-05) (Table S2, Model 1).

In the second analysis (using all subjects with the adjustment for *APOE ε4*), we identified seven independent SNPs (p<5e-07) (Fig. 1B, Table S2, Model 2), including three aforementioned variants in *TREM2*, *CLU* and *PILRA*. Four additional variants include rs144292455 in *TACR3* on 4q24 (HR=5.53, p=5.39e-07, MAC=17), rs111033333 in *USH2A* 1q41 (HR=4.58, p=2.74e-07, MAC=19), rs199533 in *NSF* on 17q21.31 (HR=0.87, p=3.95e-07, minor allele frequency (MAF)=20.2%), and rs61981931 in *CDKL1* 14q21.3 (HR=0.77, p=4.57e-07, MAF=4.9%). The SNP rs199533 in *NSF* was previously reported in [18], but did not reach the genome-wide significance in a follow-up meta-analysis incorporating replication studies [18]. The other three variants are novel. This analysis also identified three variants in *CST9* and *STAG3* genes at the suggestive level of significance p<5e-06 (Table 1).

**Table 1.** Summary statistics of candidate SNPs associated with age-of-onset of AD identified from ADSP in the analysis using all subjects adjusted for *APOE ε4* and the analysis using *APOE ε4* non-carriers. POS: coordinate of the SNP in build 37. log(HR): log of hazard ratio. Assumption test: P-values for testing the assumption of proportional hazards in ADSP.

| SNP | CH<br>R | POS | GENE | MAF<br>ADSP | ADSP |  | ROSMAP |  | CHS |  | LOADFS |  | GENADA |  | ADSP<br>EX | ADSP<br>EX | META-<br>ANALYSIS |  | ASSUMPT<br>ION TEST |
| --- | --- | --- | --- | --- | --- | --- | --- | --- | --- | --- | --- | --- | --- | --- | --- | --- | --- | --- | --- |
|  |  |  |  |  | log(H<br>R) | P | log(H<br>R) | P | log(H<br>R) | P | log(H<br>R) | P | log(H<br>R) | P | log(H<br>R) | P | log(H<br>R) | P | P |
| ANALYSIS USING ALL SUBJECTS ADJUSTED FOR APOE E4 |  |  |  |  |  |  |  |  |  |  |  |  |  |  |  |  |  |  |  |
| rs111033333 | 1 | 216270469 | USH2A | 9.30E-04 | 1.52 | 2.74E-07 | NA | NA | NA | NA | 0.22 | 6.94E-01 | 0.24 | 6.33E-01 | NA | NA | 1.03 | 9.73E-06 | 6.08E-01 |
| rs144292455 | 4 | 104577415 | TACR3 | 8.34E-04 | 1.71 | 5.39E-08 | 1.28 | 2.03E-01 | NA | NA | 1.09 | 8.78E-02 | 2.19 | 3.06E-02 | 0.34 | 7.35E-01 | 1.53 | 1.78E-09 | 1.46E-01 |
| rs1043915 | 7 | 99799845 | STAG3 | 2.52E-01 | -0.12 | 3.32E-06 | -0.02 | 8.34E-01 | -0.13 | 3.46E-01 | -0.05 | 2.77E-01 | -0.01 | 8.45E-01 | -0.03 | 6.81E-01 | -0.09 | 1.05E-05 | 5.08E-01 |
| rs61981931 | 14 | 50856882 | CDKL1 | 4.91E-02 | -0.26 | 4.57E-07 | 0.15 | 5.05E-01 | -0.06 | 8.25E-01 | -0.15 | 1.43E-01 | 0.10 | 3.56E-01 | -0.46 | 5.98E-03 | -0.19 | 2.23E-06 | 2.45E-01 |
| rs199533 | 17 | 44828931 | NSF | 2.02E-01 | -0.14 | 3.95E-07 | -0.44 | 1.82E-03 | -0.06 | 6.70E-01 | 0.07 | 1.89E-01 | -0.07 | 2.78E-01 | -0.19 | 2.20E-02 | -0.11 | 7.19E-07 | 6.34E-01 |
| rs2983640 | 20 | 23586360 | CST9 | 3.88E-01 | -0.11 | 1.42E-06 | -0.13 | 2.45E-01 | 0.06 | 6.19E-01 | -0.01 | 8.80E-01 | 0.00 | 9.96E-01 | NA | NA | -0.07 | 4.14E-05 | 4.45E-01 |
| ANALYSIS USING APOE E4 NON-CARRIERS |  |  |  |  |  |  |  |  |  |  |  |  |  |  |  |  |  |  |  |
| rs1134586 | 1 | 40980559 | EXO5 | 1.79E-01 | 0.15 | 3.08E-06 | 0.19 | 3.78E-01 | -0.39 | 1.48E-01 | 0.24 | 2.95E-02 | 0.06 | 5.93E-01 | 0.14 | 3.09E-01 | 0.14 | 4.82E-07 | 7.88E-01 |
| rs111033333 | 1 | 216270469 | USH2A | 9.30E-04 | 1.59 | 4.25E-08 | NA | NA | NA | NA | NA | NA | 0.43 | 6.68E-01 | NA | NA | 1.50 | 7.28E-08 | 5.38E-01 |
| rs144292455 | 4 | 104577415 | TACR3 | 8.34E-04 | 1.44 | 2.22E-07 | 1.26 | 2.16E-01 | NA | NA | 1.09 | 1.90E-01 | NA | NA | 0.38 | 7.02E-01 | 1.33 | 6.87E-08 | 2.27E-01 |
| rs142695278 | 9 | 105767917 | CYLC2 | 9.31E-04 | 1.45 | 1.51E-06 | NA | NA | NA | NA | -8.24 | 8.53E-01 | -10.90 | 9.92E-01 | NA | NA | 1.45 | 1.52E-06 | 7.62E-01 |
| rs149524209 | 9 | 138712860 | CAMSAP1 | 7.39E-04 | 1.53 | 1.46E-06 | NA | NA | NA | NA | NA | NA | NA | NA | NA | NA | 1.53 | 1.46E-06 | 6.85E-01 |
| rs79782048 | 9 | 139409089 | NOTCH1 | 9.80E-04 | 1.38 | 4.71E-06 | NA | NA | NA | NA | NA | NA | NA | NA | -0.67 | 5.05E-01 | 1.21 | 2.76E-05 | 6.85E-01 |
| rs12373123 | 17 | 43924073 | SPPL2C/MAPT-AS1 | 2.16E-01 | -0.14 | 5.33E-06 | -0.38 | 7.95E-02 | -0.52 | 3.61E-02 | 0.12 | 2.56E-01 | -0.23 | 3.45E-02 | -0.22 | 1.01E-01 | -0.14 | 3.55E-07 | 7.99E-01 |
| rs2732703 | 17 | 44353222 | ARL17B (intron) | 2.03E-01 | -0.14 | 1.94E-05 | -0.39 | 1.30E-01 | -0.61 | 2.24E-02 | 0.15 | 1.67E-01 | -0.20 | 7.82E-02 | -0.31 | 5.52E-02 | -0.14 | 2.74E-06 | NA |
| rs199533 | 17 | 44828931 | NSF | 2.02E-01 | -0.15 | 3.25E-06 | -0.37 | 1.08E-01 | -0.65 | 1.42E-02 | 0.13 | 2.24E-01 | -0.18 | 9.97E-02 | -0.24 | 8.33E-02 | -0.15 | 3.63E-07 | 9.58E-01 |
| rs56201815 | 17 | 62141416 | ERN1 | 1.32E-03 | 1.44 | 7.21E-08 | 1.82 | 7.40E-02 | 2.94 | 4.34E-03 | 1.86 | 1.96E-02 | NA | NA | 2.66 | 8.92E-03 | 1.64 | 2.10E-12 | 8.12E-01 |

In the third analysis (using only *APOE ε4* non-carriers), we identified four independent significant SNPs (p<5e-07) (Fig. 1C, Table S2, Model 3) including the R47H mutation in *TREM2* (HR=2.87, p=7.90e-14), rs144292455 in *TACR3* (HR=4.22, p=2.22e-07) and rs111033333 in *USH2A* (HR=4.89, p=4.25e-08) found in the second analysis. One novel SNP was the rare variant rs56201815 in *ERN1* within 17q23.3 locus (HR=4.22, p=7.21e-08, MAC=29). The HRs of the minor allele of these SNPs were substantial and comparable to that of *APOE*, which is not surprising because rare coding variants tend to show more significant biological effects, and the MAF of these SNPs in the ADSP sample is merely ~0.1%, much lower than the R47H mutation in *TREM2*. In addition, nine SNPs attaining p<5e-06 were identified (Table 1).

We found that the p-values of the newly identified SNPs from the Cox models were much more significant, particularly for the rare variants, than those from a logistic model using the same ADSP sample and covariates (Fig. 1D), explaining why these SNPs were not detected in the previous study. We compared the p-values of well-established AD-related coding-variants in the ADSP WES data between the two models. We found that the Cox model produced more significant p-values for almost all SNPs except for the two SNPs in *MS4A6A* (Fig. 1D).

### Replication analyses confirm SNPs in *ERN1, TACR3* and the *MAPT* region

The variants in *TREM2*, *CLU* and *PILRA,* identified using the full sample in our first analysis, were reported by previous larger studies [10–12]. Accordingly, we focus on replication of the novel findings identified in the analyses conditional on *APOE ε4*, and using the ε*4*-free sample. We attempted to replicate associations of 12 candidate SNPs with a p-value <5e-06 in at least one model in the discovery phase (Table 1), including six common variants (MAF≥5%) and six rare variants (MAF<1%). All these SNPs passed a test for the assumption of proportional hazards in the discovery phase (Table 1). We further included rs2732703, an intronic variant of *ARL17B* in the *MAPT* region reported being associated with AD in a previous study of *APOE ε4* non-carriers [28]. This SNP is in high linkage disequilibrium (LD) with our identified coding variants rs199533 (r^2^=0.90) in *NSF* and rs12373123 (r^2^=0.93) in *SPPL2C*. We examined these SNPs in non-Hispanic white populations of LOADFS (3473 subjects, 43.4% cases, imputed genotypes), CHS (3262 subjects, 6.2% cases, imputed genotypes), GenADA (1588 subjects, 50% cases, imputed genotypes), the Religious Orders Study (ROS) and the Rush Memory and Aging Project (MAP) cohort (1195 subjects, 45% cases, whole-genome sequencing (WGS) genotypes [29]) and the ADSP extension study (1147 subjects, 45.8% cases, WGS genotypes) (Table S1). We removed ~400 subjects from the ROSMAP WGS cohort, 572 from CHS, 318 from LOADFS, who were already included in the ADSP sample, resulting in 681, 2690, 3155 non-Hispanic whites, respectively. The coxmeg R package [21] was used to analyze the LOADFS data set with a GRM estimated from a genotype array, and the coxph function in the survival R package [30] was used to analyze the CHS, GenADA, ROSMAP and ADSP extension data sets.

Meta-analysis of the results from the conditional model adjusted for *APOE ε4* showed an improved p-value of 1.78e-09 for rs144292455 in *TACR3* (MAF=0.083% in ADSP) compared to the ADSP sample alone (p=5.39e-08). This SNP had a p-value of 6.3e-03 when a logistic model instead of a Cox model was fitted, which might be a reason for not identifying this SNP by the previous case-control study of ADSP [18]. The other SNPs did not reach the exome-wide significance of 5e-07 in the meta-analysis (Table 1). Rs144292455 is a coding variant of *TACR3* resulting in a premature stop codon and, thus a shortened transcript. The minor allele of rs144292455 increased the risk of AD in ADSP (17 carriers, 16 cases), ROSMAP (2 carriers, 1 case), LOADFS (9 subjects with a dosage>0.5), GenADA (1 subject with a dosage>0.5) and the ADSP extension study (2 carriers, 1 case), while no imputed carriers were observed in CHS (Table S7). The vast majority of the minor allele carriers in ADSP (16 of 17; 3 of 16 also carry *APOE ε4* allele) had AD with an average age-of-onset of 71.03 (Fig. 2C). This age was substantially younger than the average age-of-onset of 75.4 years based on all AD cases. Two carriers in ROSMAP were both *APOE ε4* non-carriers and the AD case carried *APOE ε2/*ε*4* genotype.

In the analysis using *APOE ε4* non-carriers, five SNPs (rs56201815, rs144292455, rs12373123, rs1134586, and rs199533) showed smaller exome-wide significant meta-analysis p-values (p<5e-07) compared to those from the ADSP sample alone. Association for rs111033333 remained at the exome-wide significance. Replication of rare variant rs111033333 was, however, less reliable as no minor allele carriers were observed in ROSMAP, LOADFS, CHS, or the ADSP extension study. The novel AD-associated SNP rs56201815 (meta-analysis p=2.10e-12) is a synonymous variant in *ERN1*. rs12373123, a missense variant of *SPPL2C* (Table 1), is located in a large LD block spanning the *MAPT region* and it is in complete LD with multiple synonymous, nonsense or missense variants in *CRHR1* and *MAPT*. In *APOE ε4* non-carriers, the hazards of AD were consistently lower in the carriers of the minor allele of rs12373123 after age 70 (Fig. 2D). It had a more significant p-value (meta-analysis p=3.55e-07) than the previously reported SNP rs2732703 (meta-analysis p=2.74e-06) and rs199533 (meta-analysis p=3.63e-07), suggesting that this SNP might be more correlated with the true causal variant for AD in this region. The minor allele of rs12373123 was consistently associated with decreased risk of AD in all studies except for LOADFS. In addition, rs1134586, a common missense variant in *EXO5*, passed the exome-wide significance with a small margin (p=4.82e-07). More data might be needed to further validate the association of this SNP.

### Minor allele of rs56201815 in *ERN1* increases the risk of AD and lowers glucose metabolism

Among the aforementioned replicated SNPs, rs56201815 in *ERN1* yielded the most significant meta-analysis p-value, and its minor allele (G) (MAF=0.15% in a non-Finnish European sample (Lek et al., 2016)) increased the risk of AD consistently across all studies and independently of the *APOE ε4* allele. The HRs were nominally significant in LOADFS (p=1.97e-02) and CHS (p=6.66e-03). In GenADA, no carriers of the minor allele were observed. We analyzed the minor allele carriers in these studies in more detail. Twenty-seven (16 males) rs56201815-G carriers in ADSP (a total of 29 carriers in which two were excluded from the analyses because they transformed from control to mild cognitive impairment (MCI) during the follow-up in ADSP, and their AD status was unknown) were sampled from 11 cohorts including ACT, ADC, CHAP, MAYO, MIA, MIR, ROSMAP, VAN, ERF, FHS, and RS (Table 2). The genotypes of these rs56201815-G carriers passed the quality control and had high sequencing depth. Of them, 23 subjects were diagnosed with AD and their average age-of-onset (73.5 years) was lower than the average age-of-onset (75.4 years) of all AD cases in ADSP (Fig. 2E). Interestingly, three of the four rs56201815-G carriers in the control group carried *APOE ε4* allele that explained why this SNP was only identified in the analysis of *APOE ε4* non-carriers. Indeed, we observed that rs56201815-G had a stronger effect on the risk of AD in *APOE ε4* non-carriers (Fig. 2F, Table S2). In the ROSMAP WGS cohort (after excluding the duplicated subjects examined in the ADSP sample), we observed three rs56201815-G carriers, including one *APOE ε4* carrier (Table 2). Two of the three carriers were diagnosed with AD, which, albeit from a small sample size, is much higher than the incidence of 36.7% in the non-carriers. The genotypes of all carriers had high sequencing quality. In the LOADFS cohort, we observed seven subjects with a dosage of rs56201815-G higher than 0.5 (Table 2). One individual was followed up until age 59 which is too young to be a useful subject for study of AD. Four out of the remaining six carriers had both AD and dementia, including three family members who were all diagnosed with AD (Table 2). This incidence (66.7%) was higher than that in non-carriers (43%). In the CHS cohort, we observed three subjects with a dosage of rs56201815-G higher than 0.5 (Table 2), one of whom (33.3%) had AD during the follow-up, higher than the incidence (6.16%) in non-carriers. In the ADSP extension WGS study, we observed two rs56201815-G carriers in non-Hispanic whites, and both were *APOE ε4* non-carriers. One of the carriers was diagnosed with AD at age 69, and the other converted to dementia during the follow-up with an unknown status of AD.

**Table 2.** Detailed information about rs56201815-G carriers in ADSP, ROSMAP, LOADFS and CHS. Age at onset/censoring: age-of-onset if the subject had AD or age at the end of follow-up if the subject was a control. AD/Non-AD: number of AD/control subjects. Male: percentage of males. APOE e4 carrier: number of *APOE ε4* carriers. Seq Quality / dosage: sequencing quality of the minor allele carriers for ADSP WES, ADSP extension WGS, and ROSMAP WGS or imputed dosage for LOADFS and CHS. Comments: additional information about the subject. Cohort: the original cohort in ADSP.

| PROJ<br>ECT | AGE AT<br>ONSET/CENS<br>ORING | TOTAL<br>SUBJEC<br>TS | A<br>D | NO<br>N-AD | MA<br>LE<br>(%) | APOE<br>E4<br>CARRIE<br>R | SEQ QUALITY<br>/ MEAN<br>DOSAGE | COMMENTS | COHORT |
| --- | --- | --- | --- | --- | --- | --- | --- | --- | --- |
| ADSP | 60-70 | 6 | 6 | 0 | 50 | 4 | PASS |  | ADC, MIR, VAN |
|  | 70-80 | 13 | 1<br>3 | 0 | 53.8<br>5 | 5 | PASS |  | ADC, MAYO, MIA,<br>MIR, ROS, VAN, ERF,<br>FHS, RS |
|  | >80 | 8 | 4 | 4 | 75 | 3 | PASS |  | ACT, ADC, CHAP, ROS,<br>RS |
|  | NA | 2 | N<br>A | NA | 50 | 0 | PASS | Control converted to MCI. Not<br>included in the analysis due to<br>unknown AD status. | ADC |
| ROSM<br>AP* | 80-90 | 3 | 2 | 1 | 0 | 1 | 99 |  |  |
| LOADF<br>S | <60 | 2 | 1 | 1 | 100 | 0 | 0.84 | The control subject was censored at age 59. |  |
|  | 60-82 | 5 | 3 | 2 | 80 | 1 | 0.856 | One of the cases was not included in the analysis due to<br>unknown exact age-at-onset. This subject also had dementia at<br>age 73. |  |
| CHS | 70-80 | 3 | 1 | 2 | 100 | 0 | 0.93 |  |  |
| ADSP<br>extensi<br>on | 60-70 | 1 | 1 | 0 | 100 | 0 | 99 |  | ADC |
|  | NA | 1 | N<br>A | NA | 100 | 0 | 99 | This subject converted to dementia<br>during follow-up. | ADC |
| * Not include the three subjects<br>overlapping in ADSP |  |  |  |  |  |  |  |  |  |

The ADNI project was not included in the replication analysis because the age-of-onset of AD was not available. Moreover, the vast majority of the ADNI WGS sample (738 subjects) was MCI or control subjects, and AD cases accounted for merely 5.8%. Instead, we investigated the association between rs56201815 and average FDG-PET intensity, one of the most accurate biomarkers to predict conversion from MCI to AD and to distinguish between control, early MCI (EMCI), late MCI (LMCI) and AD subjects [31–35], across five brain regions of interest (ROIs) (left/right angular gyrus, bilateral posterior cingulate gyrus, and left/right inferior temporal gyrus). We observed that the average FDG uptake of the five rs56201815-G carriers (two LMCI subjects, one EMCI subject, and two controls) adjusted for within-subject variability, age at measurement, sex, and diagnosis groups (control, EMCI, LMCI and AD) was significantly lower than that of the homozygous subjects (Fig. 3A), suggesting that the rs56201815-G carriers had lower cerebral glucose metabolism and will more likely convert to advanced stages.

**Figure 3.**
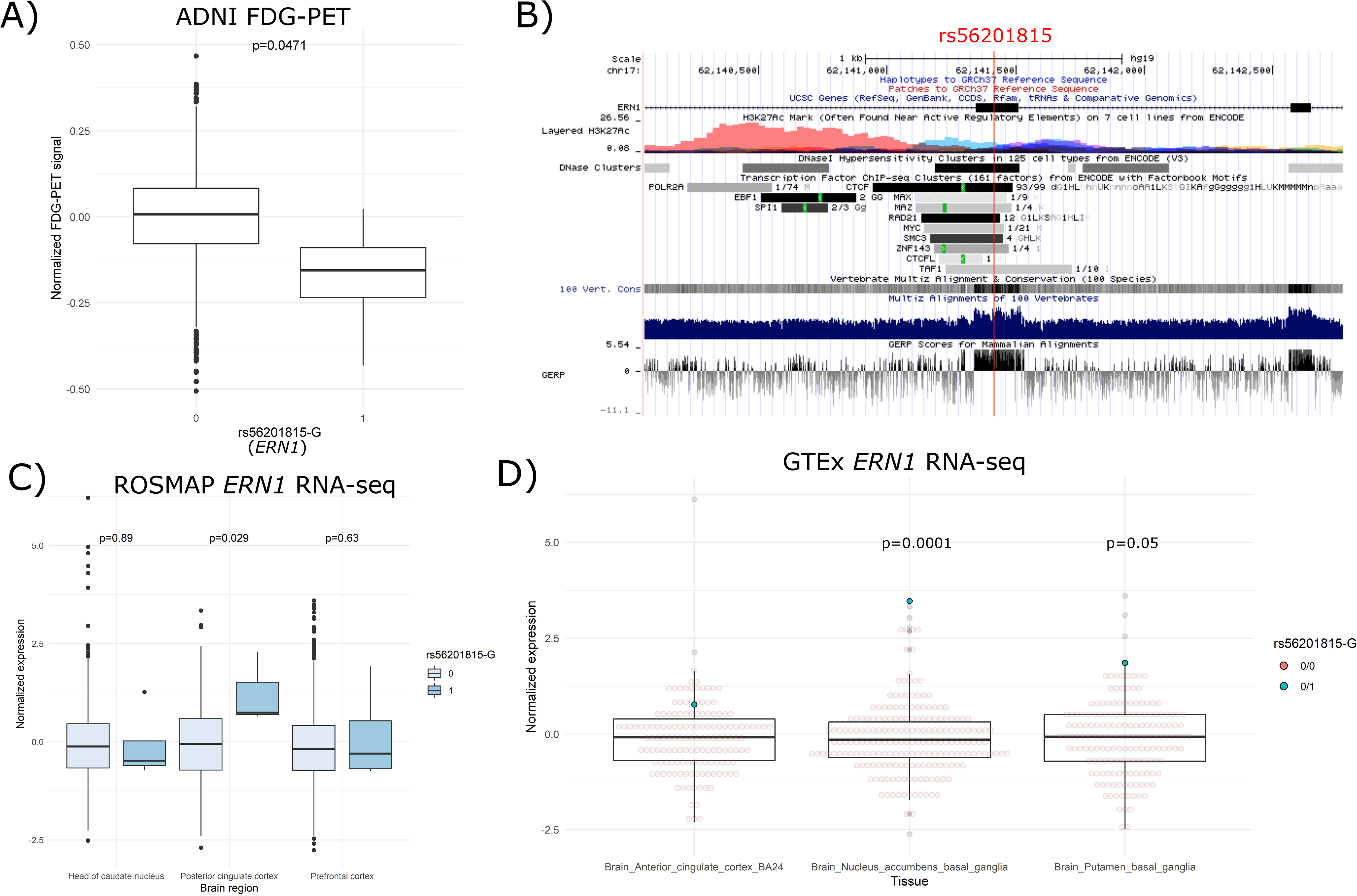
(A) Normalized longitudinal FDG-PET measurements between rs56201815-G carriers and non-carriers in ADNI. The p-value was calculated using a linear mixed-effects model in which individual-level random effects and three covariates (age at the measurement, sex and diagnosis) were adjusted. (B) Annotation of histone modifications, transcriptional factor binding, and evolutionary conservation in the genomic region of rs56201815. (C) Normalized expression of *ERN1* between rs56201815-G carriers and non-carriers in three brain tissues in cerebrum (dorsolateral PFC, PCC and anterior caudate nucleus) from a ROSMAP RNA-seq sample. (D) Normalized expression of *ERN1* between rs56201815-G carriers and non-carriers in anterior cingulate cortex, nucleus accumbens, and putamen from GTEx RNA-seq samples.

### rs56201815 is synonymous variant and brain-specific eQTL of ERN1

As rs56201815 in *ERN1* was the most significant SNP identified from the discovery and replication phases, we next sought to examine its biological and regulatory functions. rs56201815 is a synonymous coding variant, indicating that it unlikely alters amino acid sequence of *ERN1*. However, rs56201815 is located in a CTCF binding site, an open chromatin region in multiple cell types and an evolutionarily conserved region (Fig. 3B). Moreover, a recent mouse study reports that inhibition of *ERN1* expression reduces amyloid precursor protein (APP) in cortical and hippocampal areas, and restores the learning and memory capacity of AD mice [36]. We, therefore, hypothesized that rs56201815 is a cis-eQTL of *ERN1* in brain, and the detrimental effect of rs56201815 on AD is mediated by upregulating the expression of *ERN1*. To test this hypothesis, we examined the effect of rs56201815 on the expression of *ERN1* using RNA-seq data in ROSMAP and GTEx, and microarray data in ADNI.

We collected 2213 RNA-seq samples from 838 subjects in the ROSMAP cohort in three brain regions including dorsolateral prefrontal cortex (PFC), posterior cingulate cortex (PCC), and anterior caudate nucleus, among which four subjects were rs56201815-G carriers. Our differential expression (DE) analysis revealed that the minor allele of rs56201815 was associated with increased expression of *ERN1* (log(fold-change (FC))=0.204, p=0.0285) in PCC (Fig. 3C). We then analyzed a WGS data set of 838 healthy subjects from the GTEx project. The WGS data included two rs56201815-G carriers. One of them had RNA-seq data in nine brain tissues including amygdala, anterior cingulate cortex (ACC), hypothalamus, caudate, nucleus accumbens, putamen, cerebellar hemisphere, cerebellum and spinal cord. Despite the small sample size, our DE analyses indicated that rs56201815 was a potential eQTL of *ERN1* in several regions in cerebrum, particularly nucleus accumbens (log(FC)=1.28, p=1e-4), and putamen (log(FC)=0.734, p=0.05) (Fig. 3D). In line with the result from the ROSMAP data in PCC, rs56201815-G was correlated, albeit not significant (log(FC)=0.35, p=0.437), with the expression in ACC, leading to a significant meta-analysis p-value of 0.0213 for cingulate cortex. In almost all regions in cerebrum, the rs56201815-G carrier had uniformly higher expression of *ERN1* than the average (Fig. 3D, Fig. S4A).

We then investigated the effects of rs56201815 on *ERN1* expression in other brain regions, and in four non-brain tissues including sigmoid colon, lung, spleen and whole blood. The RNA-seq data in sigmoid colon had two rs56201815-G carriers, and one rs56201815-G carrier was available in the other tissues. The DE results showed no evidence of an association between rs56201815 and the gene expression in any of these tissues (Fig. S4A). As the number of rs56201815-G carriers in the GTEx project is small, we further analyzed a peripheral whole blood sample from the ADNI project, comprising 733 subjects having both a WGS dataset and a microarray gene expression dataset, three of whom were rs56201815-G carriers with high sequencing quality. Our DE analyses of two probes in *ERN1* showed that the minor allele rs56201815-G was not associated with either probe (Fig. S4B).

These results suggested that rs56201815 was associated with elevated expression of *ERN1* in cerebral regions (most predominantly in PCC and several regions in basal ganglia), but not likely in other tissues. To examine whether its regulatory effects in the brain are mediated by a change of chromatin activity, we further carried out association analyses of epigenetic markers including DNA methylation and histone modifications in PFC. We collected an Illumina 450k array DNA methylation data set of 721 subjects (four rs56201815-G carriers) from a ROSMAP sample [37, 38]. Among 11 probes located in the region of *ERN1*, we found no evidence of significant association after adjustment for multiple testing (Table S3). The most significant probe (chr17:62134117), also the probe closest to rs56201815, was located in an enhancer with a p-value of 0.012. For histone modifications, we interrogated histone 3 lysine 9 acetylation (H3K9ac) peaks using a ChIP-seq data set of 632 subjects (four rs56201815-G carriers) from a ROSMAP sample [37, 39]. We conducted differential analyses of 26,384 broad peaks adjusted for fraction of reads in peaks (FRiPs), GC bias, and 10 remove unwanted variation (RUV) components. No significant association was found among nine broad peaks within a ±200kb flanking region of *ERN1* after adjustment of multiple testing although eight peaks showed slightly increased intensity in the carriers (Table S4). The most significant association was in an enhancer at chr17:62,337,374-62,342,372 with a p-value of 0.043.

### Rs12373123 is neural cell type-specific eQTL of MAPT and GRN

Previous studies show that rs12373123 is a cis-eQTL of multiple nearby genes (e.g., *MAPT*, *CRHR1*, and *LRRC37A*) in multiple tissues including brain [28, 40–42], and shows chromatin interactions with these genes (Fig. 4A). But it is not clear which cell type and genes mediate its effect on AD. We then explored the regulatory effects of rs12373123 at a cell-type level using a single-nucleus RNA-seq (snRNA-seq) dataset. Cell type-specific analysis can also reduce the potential confounding effects originating from unobserved heterogeneous cell type proportion across subjects in the tissue-level analysis, and therefore produces more accurate and refined estimates. We performed cell type-specific eQTL analyses using 44 subjects having both genotype data (39 subjects from WGS and 5 subjects from a SNP array) and snRNA-seq data from ~80,000 cells in PFC from a ROSMAP sample. We classified cells into excitatory neurons, inhibitory neurons, astrocytes, microglia, oligodendrocytes, and oligodendrocyte progenitor cells (OPCs) based on previous clustering results [43]. We then aggregated cells within each cell type and each subject.

**Figure 4.**
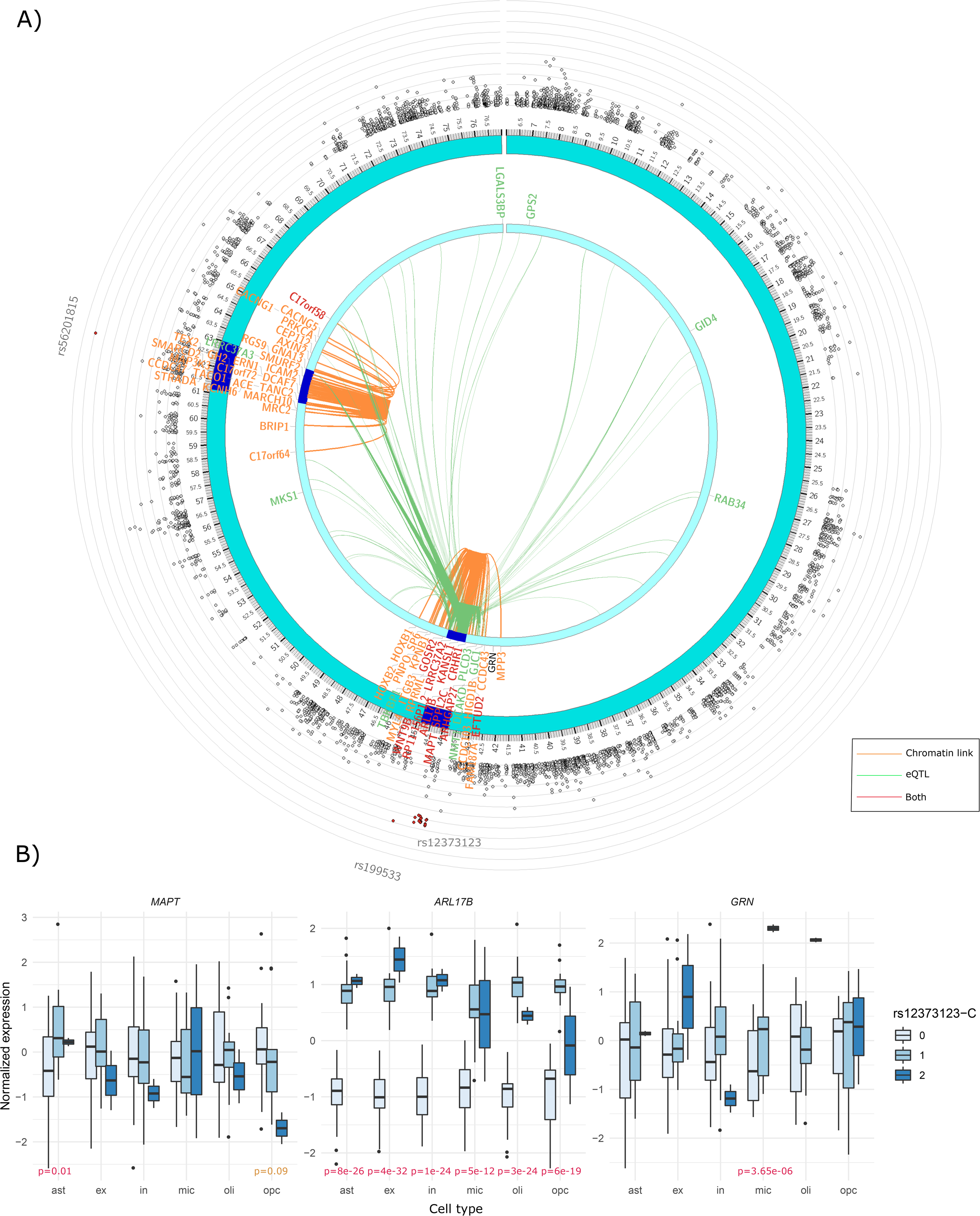
(A) Chromatin interaction (orange links) and tissue-specific eQTLs (green links) for rs56201815 and rs12373123 on chromosome 17 identified from the exome-wide association analysis of age-of-onset of AD in *APOE ε4* non-carriers in ADSP. A gene that is in chromatin interaction or an eGene with these SNPs is highlighted in orange or green, respective. A gene highlighted in red indicates both features. (B) Normalized expression of three genes (*MAPT, ARL17B,* and *GRN*) near rs12373123 of 44 subjects (including 13 rs12373123-T/C carriers and 2 rs12373123-C/C carriers) in six major brain cell types (astrocytes, excitatory neurons, inhibitory neurons, microglia, OPCs and oligodendrocytes). All cells in each cell type from each subject were first pooled, and the gene expression were aggregated by subjects. The gene expression was then adjusted for age, sex, and AD status.

In each cell type, we interrogated 11 protein-coding genes (10 genes within a ±500kb flanking region and *GRN*, a nearby gene linked to frontotemporal lobar degeneration (FTD), a type of dementia). The cell type-specific eQTL analyses revealed that one or more copies of rs12373123-C were associated with elevated expression of *ARL17B* in all six brain cell types (p<1e-11) (Fig. 4B, Table S5). rs12373123 was also an eQTL of *LRRC37A3*, *KANSL1*, and *CRHR1* in all cell types except for microglia (Fig. S2, Table S5). The protective allele rs12373123-C was associated with elevated *MAPT* expression in astrocytes (p=0.01) while a decreasing trend in OPCs (p=0.09) (Fig. 4B, Table S5). We further found that rs12373123-C, particularly its homozygous protective genotype, was significantly associated with increased expression of *GRN* in microglia (p=3.65e-06) (Fig. 4B, Table S5), which is a protective gene against dementia and is important for lysosome homeostasis in the brain [44, 45].

We also assessed the cell type-specific association between rs56201815 and the expression of *ERN1*. We observed that *ERN1* was ubiquitously expressed in all brain cell types, most abundantly in microglia, followed by astrocytes and OPCs. As there was only one rs56201815-G carrier among the 39 WGS subjects, and, unfortunately, its total sequencing depth was much lower than that of the other subjects (~10% of the average library size), we investigated three major abundant cell types (excitatory neurons, astrocytes, and oligodendrocytes), for which the carrier had a library size > 50,000. We observed that rs56201815-G was slightly correlated with increased expression of *ERN1* in excitatory neurons, but not significant (Fig. S3).

### Gene-set analysis identifies astrocyte, microglia and amyloid-beta related pathways

As aggregating signals within a gene can often increase the statistical power, in particular, for detecting rare coding variants, we carried out gene-based analyses using the summary statistics of all examined SNPs estimated from the ADSP sample. Our gene-based analyses using MAGMA [46] showed that *TREM2* was the most significant gene associated with AD in all individuals (p=1.65e-9) and in *APOE ε4* non-carriers (p=2.57e-10) (Fig. S5A), consistent with previous results [18]. Indeed, all six exonic SNPs (rs2234256, rs2234255, rs2234253, rs142232675, rs143332484, rs75932628) in *TREM2* were at least nominally associated with AD (Table S2). Its significance in *APOE ε4* non-carriers was higher, suggesting that the effects of *TREM2* on AD were independent of *APOE*. Besides, multiple genes in the *MAPT* region including *MAPT, KANSL1, NSF,* and *SPPL2C* were significantly (p<3.5e-06) associated with the risk of AD in both analyses (Fig. S5B). We also observed that *CLU, TACR3, NAV2,* and *FAM186B* were among the top associated genes.

Our gene-set analysis using FUMA [47] based on the summary statistics from the exome-wide association analysis conditional on *APOE ε4* detected one Gene Ontology (GO) gene set (positive regulation of astrocyte activation) associated with AD (Bonferroni adjusted p<0.05) (Fig. 5A). Other top enriched gene sets included antigen processing and presentation via MHC class II, amyloid-beta clearance and formation, response to endoplasmic reticulum (ER) stress, and microglial activation (Fig. 5A). Consistently, the gene sets related to astrocytes, microglia, and amyloid-beta were also among the top in the gene-set analysis using *APOE ε4* non-carriers (Fig. 5B). Our cell type association analysis using FUMA (Watanabe et al., 2019) showed that microglia were associated with AD among nine major cell types in the brain (<0.05) in the analysis of *APOE ε4* non-carriers (Fig. 5D). No cell type was associated with AD based on the summary statistics from the association analysis conditional on *APOE ε4* (Fig. 5C).

**Figure 5.**
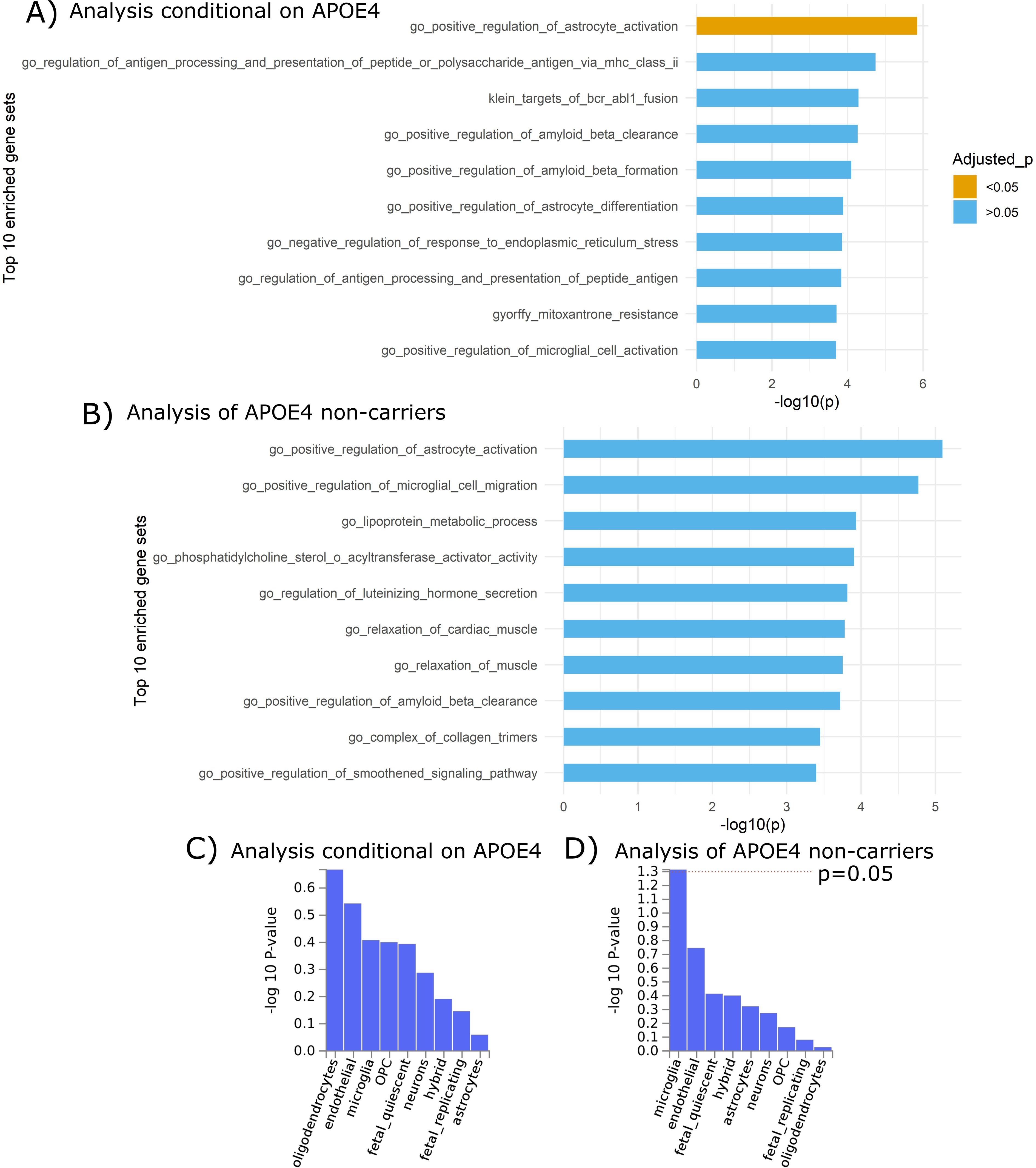
Top 10 gene sets enriched based on the exome-wide association analyses of age-of-onset of AD using (A) Model 2: a model with all subjects adjusted for copy of *APOE ε4*, PC2 and sex; (B) Model 3: a model with *APOE ε4* non-carriers adjusted for PC2 and sex. Brain cell-type enrichment analysis based on the exome-wide association analyses of age-of-onset of AD using (C) Model 2 and (D) Model 3.

## Discussion

In this study, we interrogated the associations between 108,509 exome-wide SNPs and age-of-onset of late-onset AD using Cox models with a sample consisting of ~20,000 AD patients and controls. We also attempted to identify SNPs contributing to earlier onset in *APOE ε4* non-carriers alone. Most of these SNPs are rare variants. Our results not only confirm previously reported AD-related SNPs with much higher significance but also reveal novel genetic variants associated with age-of-onset of AD, particularly in *APOE ε4* non-carriers.

One of our major findings is a synonymous rare variant, rs56201815, in *ERN1* (also known as *IRE1*). Our results showed that the minor allele of this SNP was associated with a dramatically higher risk of AD, particularly in *APOE ε4* non-carriers. Its huge effect size unanimously replicated in three other cohorts, is not surprising as its MAF in the population is only ~10% of the rare variant rs75932628 in *TREM2* according to ExAC (https://gnomad.broadinstitute.org/). *ERN1* encodes a key protein, containing a serine/threonine-protein kinase domain and a ribonuclease (RNase) domain, involved in unfolded protein response (UPR) to ER stress by activating its downstream target *XBP1* [48, 49]. Interestingly, a recent experimental study shows that the proportion of activated *ERN1* in postmortem brain tissue is associated with a Braak stage of advanced AD patients [36]. Deactivation of the RNase domain of *ERN1* in neurons reduces all hallmarks of AD including amyloid-beta load, cognitive impairment, and astrogliosis in 5xFAD mice [36]. Moreover, the ablation of eIF2α kinase PERK, one of the three major UPR genes, also prevents defects in synaptic plasticity and spatial memory in AD mice [50]. Our findings show that the minor allele of rs56201815, increasing mRNA expression of *ERN1* in multiple brain regions, also significantly increases the risk of AD, which corroborate these experimental results and provide more evidence that responses to ER stress are probably involved in the causal pathway of AD.

Aging is the most important risk factor for late-onset AD, indicating that certain risk factors during the aging process might be implicated and required in the pathogenesis of AD. The UPR is one of the mechanisms disrupted during aging, resulting in augmented susceptibility to ER stress and the accumulation of unfolded protein [51]. Previous studies show that aging leads to deficits in the systems involved in the defense against unfolded proteins in the rat hippocampus [52]. Persistent ER stress in the central nervous system during aging can initiate apoptosis of neurons and can trigger the innate immune response in microglia [53, 54]. Combined with the fact that many AD-related genes identified by GWAS are expressed exclusively in microglia, our findings indicate that the interaction between the UPR and innate immune system might play a critical role in biological mechanisms underlying AD.

As rs56201815, the variant rs12373123 in the *MAPT* region was also identified in *APOE ε4* non-carriers. The minor allele of rs12373123 was associated with reduced susceptibility to AD in ADSP, ROSMAP, CHS, and GenADA. This SNP is located in an LD block spanning >400kb, and is in high LD with a large number of SNPs including multiple missense variants in *MAPT*, *SPPL2C*, *CRHR1* and *KANSL1*. Previous GWAS show that rs12373123 and two nearby missense SNPs (rs12185268 and rs12373124) in complete LD with rs12373123 exhibit pleiotropic associations with numerous diseases and traits including intracranial volume [55], corticobasal degeneration [56], PD [57–60], primary biliary cirrhosis [61], red blood cell count [62], and androgenetic alopecia [63]. On the other hand, the major allele, more predisposed to degenerative diseases, is significantly associated with increased bone mineral density [64, 65]. Because SNPs contributing to age-related degenerative diseases are generally not subject to evolutionary selection [66, 67], its major allele is probably selected by evolution due to its beneficial effect on bone mineral density. The results of our age-of-onset analyses indicate that this pleiotropic region might also be implicated in late-onset AD, especially in *APOE ε4* non-carriers. Our cell type-specific analyses reveal that rs12373123 is a cis-eQTL in different brain cells of multiple critical genes implicated in PD and FTD (e.g., *MAPT* and *GRN*), elucidating the regulatory mechanisms underlying its pleiotropy. Due to the involvement of tau protein in the etiology of AD and PD, the effect of rs12373123 on these diseases might be mediated by *MAPT*. Indeed, rs12373123 is in high LD with multiple missense SNPs (e.g., rs62056781 and rs74496580) in *MAPT*, and we found in the snRNA-seq data that rs12373123 is also an eQTL of *MAPT* in astrocytes. Our finding also suggests that the effects of rs12373123 can be mediated by increasing the expression of *GRN* in microglia, which is a key gene protective against FTD.

*TACR3* encodes NK3-R, one of the neuropeptide neurokinin receptors that is predominantly expressed in most brain regions [41]. Stimulation of NK3-R significantly enhances learning and memory performance in aged rats [68]. Furthermore, a SNP located in 3’-UTR of *TACR3* is associated with learning performance, hippocampal volume in elderly human patients [68], and the most anterior basal forebrain volume [69]. The association of rs144292455, a nonsense variant in *TACR3,* provides more evidence for its potential involvement in AD.

Also, our results demonstrated advantages in the statistical power of using a Cox model for age-of-onset traits than a logistic model for binary outcomes in the study of AD. The power gain in terms of p-values is evident for many well-known AD-related SNPs in e.g., *TREM2* and *CLU*, which all achieved more significant p-values than a previous study using the same cohort [18]. Despite a smaller sample size, the p-value from the Cox model for detecting *APOE ε4*, the recognized true positive signal, is much more significant than a recent large-scale meta-analysis of AD status [10] and a previous analysis using a linear model of log-transformed age-of-onset [26]. Moreover, our age-of-onset analysis showed promising results for identifying rare variants compared to logistic regression. An advantage of a Cox model over Poisson regression or logistic regression is that it implicitly accounts for age-varying hazards, a characteristic in many age-related diseases, e.g., AD [70]. Our results in AD suggest that Cox models can have a power advantage for exploring rare variant association in other age-related diseases.

Although our identified SNPs were validated in multiple independent cohorts, we acknowledge some limitations. The definitions and criteria of diagnosis of AD can vary across these cohorts. AD has a certain similarity in the clinical and biological manifestation of other common neurodegenerative diseases such as FTD, which makes the clinical diagnosis of AD more complicated. Also, two of our findings rs56201815 in *ERN1* and rs144292455 in *TACR3* are rare variants (MAF=~0.13% and ~0.09%), which had lower imputation quality compared to a common variant, especially for rs144292455. Most of our GWAS replication cohorts had moderate imputation quality for rs56201815. Although these SNPs showed solid associations in our meta-analyses, as the sample sizes of our WGS replication cohorts are small for rare variants, more GWAS studies using large-scale WGS or WES data are preferable to further validate these identified associations.

In conclusion, we identified three novel SNPs in *ERN1, TACR3,* and *SPPL2C/MAPT-AS1* that exhibit strong associations with the age-of-onset of AD. We also explored their regulatory consequences at the tissue and single-cell levels in the brain. These findings support the hypothesis of the potential involvement of the UPR to ER stress and tau protein in the pathological pathway of AD, contributing to the understanding of the biological mechanisms underlying AD. Our findings are useful for guiding follow-up studies and provide more insight into the molecular mechanisms and implications of the relevant genes in AD.

## Methods

### Phenotypes in age-of-onset GWAS

A total of 10,913 European-American participants used in the discovery phase of the exome-wide age-of-onset association analyses of AD were collected from the ADSP project. These subjects were sampled from 24 cohorts, among which >3000 subjects were sampled from the ADC project (Table S6). The AD status of individuals used in the analyses was defined by clinical assessment based on NINCDS-ADRDA criteria of AD. All controls were cognitively normal individuals aged 60+. Details about study design and sample selection were described in [71]. The AD status variable in the ADSP dataset was constructed based on information on prevalent and incident AD status from the updated dataset (Version 7 with release date on June 09, 2016) if available. Otherwise, information on prevalent and incident AD status as given in Version 5 (release data on July 13, 2015) was used. More specifically, a subject was treated as AD if either prevalent or incident AD status during the ADSP follow-up was observed. The age-of-onset variable was based on the same datasets as the AD status. In both versions (Version 5 and 7), all data for age-of-onset, which we received from dbGaP, were censored by age 90.

Six cohorts (ROSMAP, LOADFS, CHS, GenADA, the ADSP extension study) were included in the replication phase of the age-of-onset GWAS. To be consistent with the AD status in ADSP, AD status in ROSMAP was based on the clinical diagnosis of AD at the last visit. For AD cases, the age at first Alzheimer’s dementia diagnosis variable was used as age-of-onset, which was also censored by age 90 if it was 90+. For controls, age-of-onset was calculated as age at the last visit or age at death if age at the last visit was not available. In LOADFS, some subjects had missing information about the age-of-onset of AD. For these subjects, we treated them as censored and set its age-of-onset as the age at the recruitment. In CHS and GenADA, the AD status and age-of-onset variables in phenotype files provided in dbGaP were used. In the ADSP extension study, the ‘AD’ and ‘Age’ variables in phenotype files were used as the AD status and the age-of-onset. We included definitive AD and control subjects, and subjects diagnosed with probable AD, possible AD, family AD, non-family AD, or unknown were not included in the analysis.

### Genotyping, Imputation, and Quality Control

WES genotypes of bi-allelic SNPs mapped to hg19 from 10,913 ADSP participants were called using the quality-controlled Atlas-only pipeline at Baylor College of Medicine (We did not use the data from the GATK pipeline at the Broad institute due to known quality issues (https://www.niagads.org/adsp/data-notices)). More details about the production of the WES data in ADSP can be found in [18]. Variants with missing rate > 2% or MAC < 10 were excluded from the age-of-onset association analyses. After the filtering, 108,509 and 110591 variants remained in the analysis using all subjects and *APOE ε4* non-carriers, respectively. VCF files of recalibrated WGS data from 1196 participants in ROSMAP were downloaded from the synapse website (https://www.synapse.org/). A total of 681 subjects were included in the replication phase after removing 16 discordant WGS samples, 17 duplicates and 477 subjects overlapping the ADSP sample. Genotyping of 3043 participants in CHS was performed using a HumanOmni1-Quad Illumina array. Genotyping of 3456 non-Hispanic Caucasian participants in NIA-LOADFS was performed using a Human610-Quad Illumina array (~600K SNPs).

Genotyping of 1588 non-Hispanic Caucasian participants in GenADA was performed using two Affymetrix 250K arrays (a total of ~500K SNPs). More information about these cohorts can be found in [72–74]. We phased and imputed the genotypes in these cohorts using the Michigan imputation server [75] with a reference panel from the Haplotype Reference Consortium (HRC) (Version r1.1 2016). WGS project level genotype VCF files (hg38) called by GATK were downloaded from NIAGADS (https://dss.niagads.org/datasets/ng00067/), from which the genotypes of 1147 non-Hispanic whites in the ADSP extension study were extracted.

### Exome-wide age-of-onset association analysis

The association analyses of the age-of-onset of AD in the discovery phase of ADSP was conducted using a Cox mixed-effects model implemented in the coxmeg R package [21], which accounted for the family structure in the cohort using a genetic relatedness matrix (GRM). A dense GRM was first estimated from the original WES data based on the GCTA model [76] implemented in the SNPRelate R package [77]. In the discovery phase of ADSP, we built a sparse GRM by setting any entry below 0.03 to zero. We evaluated five top PCs (PC1 to PC5) calculated from the dense GRM, and included the only significant PC2 in the analyses. We first estimated a variance component in the null model, which was then used to estimated HRs and p-values for all SNPs. We performed two analyses, (a) including all subjects with PC2, sex and the number of copies of *APOE ε4* included as covariates, (b) including only *APOE ε4* non-carriers with PC2 and sex included as covariates. We found that the estimated variance component was zero in the analysis (b), suggesting no evidence of random effects, and therefore we instead used a simple Cox model. The threshold to declare significant associations was calculated as 0.05 divided by the total number of tested SNPs. For comparison with the analysis of AD status, we performed association analysis by fitting a logistic regression using the glm R function adjusting for the same covariates with the same sample.

We performed age-of-onset association analyses in LOADFS, CHS, ROSMAP, GenADA, and the ADSP extension study for the top SNPs passing the suggestive threshold (p<5e-06) in the discovery phase. The same model and estimation procedures as in ADSP were used in LOADFS, which is also a family-based cohort. In LOADFS, the GRM was estimated from the genotype array data. The association analyses were conducted in the other four cohorts (i.e., CHS, ROSMAP, GenADA and the ADSP extension study) using a Cox model implemented in the survival R package [30] because these cohorts consisted of unrelated subjects. We also included sex and the number of copies of *APOE ε4* as covariates. Meta-analysis effect sizes and standard errors were computed using the summary statistics from all five studies based on the following fixed-effects model,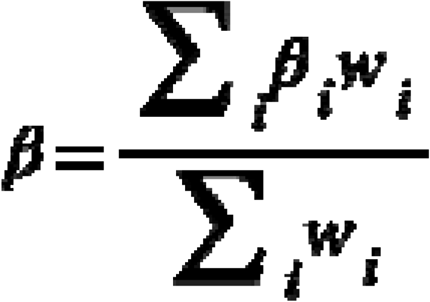 and 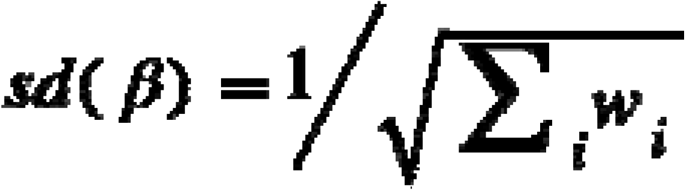, where is the weight for the study. To compare age-of-onset analysis with case-control analysis, we also performed association analyses of AD status in ADSP using logistic regression.

### Gene-based association analysis

The gene-based analysis was performed based on the summary statistics obtained from the age-of-onset association analyses. We only included SNPs with MAC≥10 and missing rate<2% in the gene-based analyses. Each SNP was first annotated to a gene using its SNP ID according to a gene location file obtained in the MAGMA website (https://ctg.cncr.nl/software/magma). We only included SNPs within the boundary of a gene body. Gene-based p-values were then computed using MAGMA with a SNP-wise mean model [46]. LD between the SNPs was estimated using the raw WES data in ADSP.

### Gene-set and cell type association analysis

The gene-set analysis was performed for curated gene sets and GO terms using the procedure SNP2GENE in FUMA [47] based on the summary statistics obtained from the age-of-onset association analyses. The 1000 Genomes Project (phase 3) for the European population was used as a reference panel in the analysis. The cell type association analysis was also performed using FUMA [78] following the SNP2GENE procedure. We selected a human brain single-cell RNA-seq data set provided in [79] as a reference for cell type-specific gene expression.

### Analysis of FDG-PET data

The longitudinal FDG-PET average intensity scores across five regions of interest (ROIs) (left/right angular gyrus, bilateral posterior cingulate gyrus, and left/right inferior temporal gyrus) for 738 subjects in ADNI having the WGS data were downloaded from the ADNI website (https://ida.loni.usc.edu). Details about sample preparation and data generation were described in [32, 33]. The association analysis between average FDG-ROI and the genotype of rs56201815 was performed by fitting a linear mixed-effects model using lme4 R package [80] including a random effect accounting for within-subject variability and three covariates (age, sex, and diagnosis group).

### Analysis of tissue-specific RNA-seq and microarray data

BAM files of aligned reads from a total of 2213 RNA-seq samples in three brain regions (dorsolateral PFC, PCC, and anterior caudate nucleus) in the ROSMAP project were downloaded from the synapse website (https://www.synapse.org/). Raw counts of 57,905 coding and non-coding genes were called using featureCounts [81] according to the GENCODE annotations GRCh37(r87). Samples with RIN<5 were excluded before the analysis. We first removed low-expressed genes (those genes for which fewer than three individuals had counts-per-million>1) before normalization. We then normalized the RNA-seq raw counts using the trimmed mean of M-values (TMM) normalization method [82]. In the analysis of PFC, 761 non-Hispanic Caucasian subjects (including four rs56201815-G carriers) having both gene expression and genotype of rs56201815 from the WGS data with RIN≥4.5 were included.

Differential eQTL analysis was performed using DESeq2 [83] adjusted for RIN, age at death, sex, AD status, and RNA extraction methods (polyA selection or rRNA depletion). In the analysis of PCC and anterior caudate nucleus, 371 (including three rs56201815-G carriers) and 585 (including four rs56201815-G carriers) non-Hispanic Caucasian subjects having both genotypes and gene expression with RIN≥4.5 and rRNA depletion were included, respectively.

To minimize technical noise resulted from sample preparation, we did not include polyA selection samples (accounting for merely 10% and 15% of all samples) because different RNA extraction methods have a large impact on measured expression in postmortem samples [84], and the samples of all rs56201815-G carriers were generated using rRNA depletion. Differential eQTL analysis was performed using DESeq2 adjusted for RIN, age at death, sex, AD status.

The raw count data of 3252 RNA-seq samples in nine brain tissues (i.e., amygdala, ACC, hypothalamus, caudate (basal ganglia), nucleus accumbens (basal ganglia), putamen (basal ganglia), cerebellar hemisphere, cerebellum and spinal cord (cervical c1)) and four non-brain tissues (i.e., sigmoid colon, lung, spleen and whole blood) from the GTEx project (version 8) were downloaded from the GTEx portal (https://gtexportal.org/home/datasets). Gene-level quantification was conducted by RSEM [85]. All GTEx raw count data were normalized using the same pipeline as in the analysis of ROSMAP. Differential eQTL analysis was then performed using DESeq2 with age, sex, and RIN as adjusted covariates.

The gene expression microarray data in peripheral blood from 742 ADNI subjects were profiled using the Affymetrix Human Genome U219 Array. Raw expression values were pre-processed using the robust multiarray average normalization method. More details about sample collection and data pre-processing can be found in [86]. Differential gene expression analyses were performed using linear regression adjusted for RIN and plate number.

### Analysis of DNA methylation data

The DNA methylation data in PFC were collected from 740 individuals in ROSMAP using the Illumina HumanMethylation450 BeadChip. Eighteen samples lying beyond ±3 standard deviations for the top 3 PCs were removed as outliers. We converted methylation beta-value to M-value using a logistic transformation. Differential methylation analysis was carried out using a linear regression adjusted for the top 10 PCs.

### Analysis of H3K9ac ChIP-seq data

H3K9ac ChIP-seq raw count data were downloaded from the synapse website (https://www.synapse.org/). This dataset is previously described in detail in [39]. Briefly, the sample comprising 26,384 H3K9ac peaks (nine peaks in the *ERN1* region) across the genome was collected from dorsolateral PFC of 669 subjects from the ROSMAP project, among which 625 subjects had also the WGS genotype data of rs56201815. The raw count data was normalized using the TMM method [82]. Estimation of tagwise overdispersion and the analysis of differential peaks for rs56201815 were carried out using DESeq2 [83] adjusted for FRiPs and GC bias. A sensitivity analysis was performed by further adjusting for 10 RUV components estimated using RUVSeq [87].

### Analysis of single-nucleus RNA-seq data

We collected snRNA-seq raw count data generated by [43] using the 10X Genomics Cell ranger pipeline in human PFC from 48 subjects (50% AD cases) including 17,926 genes profiled in 75,060 nuclei. We assigned cell identity and divided all cells into six subtypes (excitatory neurons, inhibitory neurons, astrocytes, oligodendrocytes, microglia, and OPCs) according to the previous clustering results [43] using the scanpy package [88]. The clustering of the cells is described in more detail in [43]. We excluded endothelial cells or pericytes because of the lack of abundant cell counts in these two cell types.

To perform cell type-specific eQTL analysis, we first merged cells in each cell type and in each subject to obtain a raw count matrix of 17,926 genes and 39 subjects (Six subjects were excluded due to lack of WGS data). We then followed the preprocessing and normalization procedures in the previous eQTL analysis of the bulk RNA-seq data. Differential eQTL analyses were then performed using DESeq2 [83] with age, sex, and AD status as covariates. RIN was not available for most of the subjects.

### Functional annotation

The epigenetic and regulatory annotation of the identified SNPs and its nearby SNPs in high LD (r^2^>0.8) was performed using Haploreg v4 [89], in which its tissue-specific epigenetic markers (H3K27ac), regulatory regions (enhancers and promoters), motif changes and eQTL information were annotated based on the ENCODE [90], Roadmap [91] and GTEx [41] projects. GWAS catalog [92] and GRASP [93] were used to annotate whether a SNP is an existing QTL.

## Acknowledgements

This manuscript was prepared using limited access datasets obtained through dbGaP (accession numbers: phs000168.v2.p2 (LOADFS), phs000572.v7.p4 (ADSP), phs000287.v3.p1 (CHS), phs000219.v1.p1 (GenADA), phs00424.v5.p1 (GTEx), NG00067.v2 (the ADSP extension study)).

This research was supported by Grants from the National Institutes of Health P01 AG043352, R01 AG047310 and R01 AG061853 to A.M.K. and RF1 AG054012, R01 AG058002, R01 AG062335, RF1 AG062377, U01 NS110453 to M.K., P30 AG10161, R01 AG15819, R01 AG17917, R01 AG36042, and R01 AG61356 to D.A.B.

The funders had no role in study design, data collection and analysis, decision to publish, or manuscript preparation. The content is solely the responsibility of the authors and does not necessarily represent the official views of the National Institutes of Health.

We are grateful to the participants in the Religious Order Study, the Rush Memory and Aging Project. Data can be requested at www.radc.rush.edu.

Data used in the preparation of this article were obtained from the Alzheimer’s Disease Neuroimaging Initiative (ADNI) database (adni.loni.usc.edu). The ADNI was launched in 2003 as a public-private partnership, led by Principal Investigator Michael W. Weiner, MD. The primary goal of ADNI has been to test whether serial magnetic resonance imaging (MRI), positron emission tomography (PET), other biological markers, and clinical and neuropsychological assessment can be combined to measure the progression of MCI and early AD. For up-to-date information, see www.adni-info.org.

See also further acknowledgements in supplementary materials Text S1.

## Author contributions

LH conceived the study. LH and YL imputed the genotype data. LH performed age-of-onset association analyses, gene-based analyses, microarray, RNA-seq, snRNA-seq, and ChIP-seq analyses. YP analyzed the DNA methylation data. DAB generated WGS, DNA methylation, H3K9Ac, and RNA-seq data in ROSMAP. AK and MK contributed to acquiring the data, and discussion of final results. All authors contributed to the writing of the manuscript.

## Supplementary Text

Text S1. Additional acknowledgements for ADSP, ADNI, GTEx, and CHS.

## Supplementary Tables

Table S1. Basic characteristics of the six study samples (ADSP, LOADFS, ROSMAP, CHS, GenADA, the ADSP extension study, and ADNI) included in the discovery and replication phases of the age-of-onset association analyses of AD.

Table S2. Summary statistics of the three age-of-onset association analyses of AD using the ADSP sample. Model 1: a model with all subjects adjusted for a PC and sex; Model 2: a model with all subjects adjusted for copy of *APOE ε4*, PC2 and sex; Model 3: a model with *APOE ε4* non-carriers adjusted for PC2 and sex.

Table S3. Summary statistics of the differential DNA methylation analyses of 11 probes in the *ERN1* region using a ROSMAP sample for rs56201815. Beta: the effect size in M-value with respect to a copy of rs56201815-G.

Table S4. Summary statistics of the differential analyses of nine H3K9ac peaks in the *ERN1* region (**±** 200k flanking region of rs56201815) using a ROSMAP sample for rs56201815. logFC: log(fold-change) with respect to a copy of rs56201815-G. logCPM: log(count per million) of the peak. LR: likelihood ratio test statistics. Classification: functional annotation of the peak. Median Count: median count of the reads in the peak across the subjects.

Table S5. Results of cell type-specific eQTL analyses of rs12373123 in six major brain cell types (excitatory neurons, inhibitory neurons, astrocytes, microglia, oligodendrocytes and OPCs) using the ROSMAP snRNA-seq data. logFC: log(fold-change) with respect to a copy of rs12373123-C. logCPM: log(count per million) of the peak. LR: likelihood ratio test statistics. Length: length of the gene body calculated by the distance between the start and end positions based on the Ensembl database “hsapiens_gene_ensembl”.

Table S6. Frequency across 24 cohorts among 10,913 subjects included in the ADSP sample.

Table S7. Detailed information about rs144292455-T carriers in ADSP, ROSMAP, LOADFS, GenADA and the ADSP extension study. Age at onset/censoring: age-of-onset if the subject had AD or age at the end of follow-up if the subject was a control. AD/Non-AD: number of AD/control subjects. Male: percentage of males. APOE e4 carrier: number of *APOE ε4* carriers. Seq Quality / dosage: sequencing quality for ADSP WES, ADSP extension WGS, and ROSMAP WGS or imputed dosage for LOADFS and GenADA. Comments: additional information about the subject. Cohort: the original cohort in ADSP.

## Supplementary Figures

Figure S1. Q-Q plots of the p-values from the exome-wide age-of-onset association analyses of AD using A) a model with all subjects adjusted for a PC and sex; B) a model with all subjects adjusted for copy of *APOE ε4*, PC2 and sex; C) a model with *APOE ε4* non-carriers adjusted for PC2 and sex. λ: genomic inflation factor.

Figure S2. Normalized expression of nine genes (*ARL17A*, *KANSL1*, *LRRC37A3*, *CRHR1*, *SPPL2C*, *LRRC37A*, *LRRC37A2, PLEKHM1*, *ARHGAP27*) near rs12373123 of 44 subjects in six major brain cell types (astrocytes, excitatory neurons, inhibitory neurons, microglia, OPCs and oligodendrocytes). All cells in each cell type from each subject were first pooled, and the gene expression were aggregated. The gene expression was then adjusted for age, sex, and AD status. Expression of *SPPL2C* was observed only in neuronal cells.

Figure S3. Normalized expression of *ERN1* of 39 WGS subjects (including one rs56201815-G carrier) in astrocytes, excitatory neurons and oligodendrocytes. All cells in each cell type from each subject were first pooled, and the gene expression were aggregated. The gene expression was then adjusted for age, sex, and AD status.

Figure S4. Normalized expression of *ERN1* between rs56201815-G carriers and non-carriers in A) ten tissues from GTEx RNA-seq samples; B) peripheral blood from an ADNI microarray sample. In the GTEx samples, there is one rs56201815-G carrier in all tissues except for colon, in which there are two carriers.

Figure S5. Results of gene-based association analyses of age-of-onset of AD in the ADSP sample using summary statistics from (A) Model 2: a model with all subjects adjusted for copy of *APOE ε4*, PC2 and sex; (B) Model 3: a model with *APOE ε4* non-carriers adjusted for PC2 and sex. Top genes with a p-value <1e-04 were highlighted. The red horizontal line is a threshold based on the Bonferroni correction (0.05/17,000=3e-06).

