## Supplementary figures and images for "Exome-wide age-of-onset analysis reveals exonic variants in *ERN1, TACR3* and *SPPL2C* associated with Alzheimer’s disease"

### Figure S1

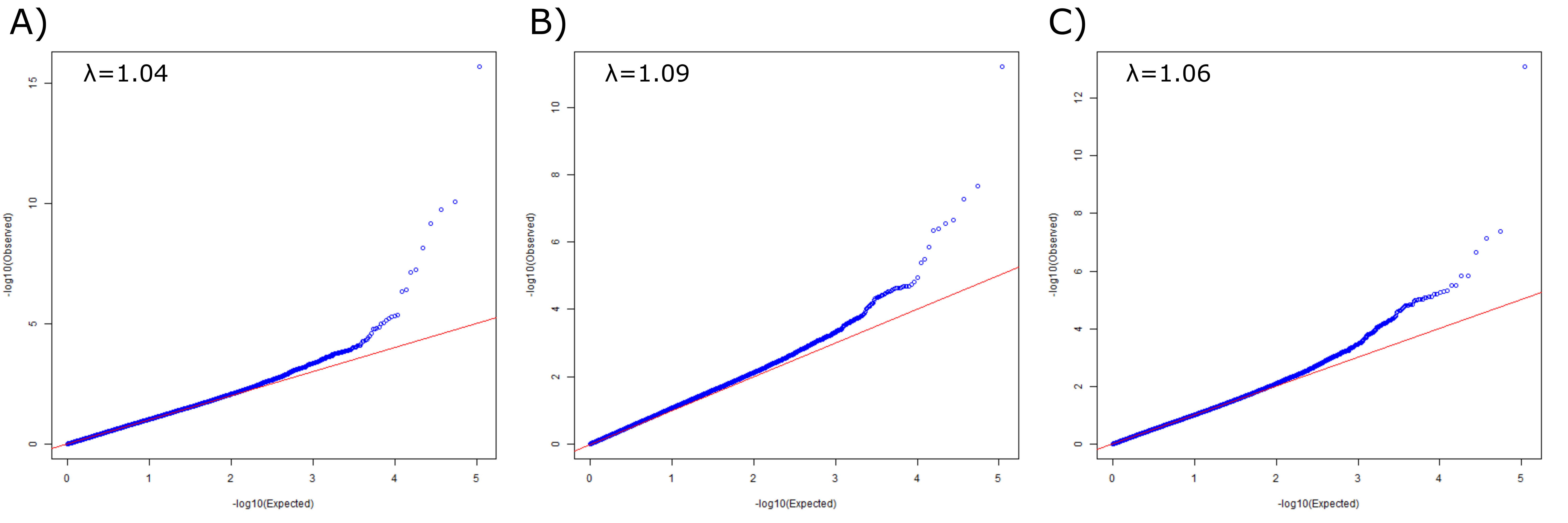

### Figure S2

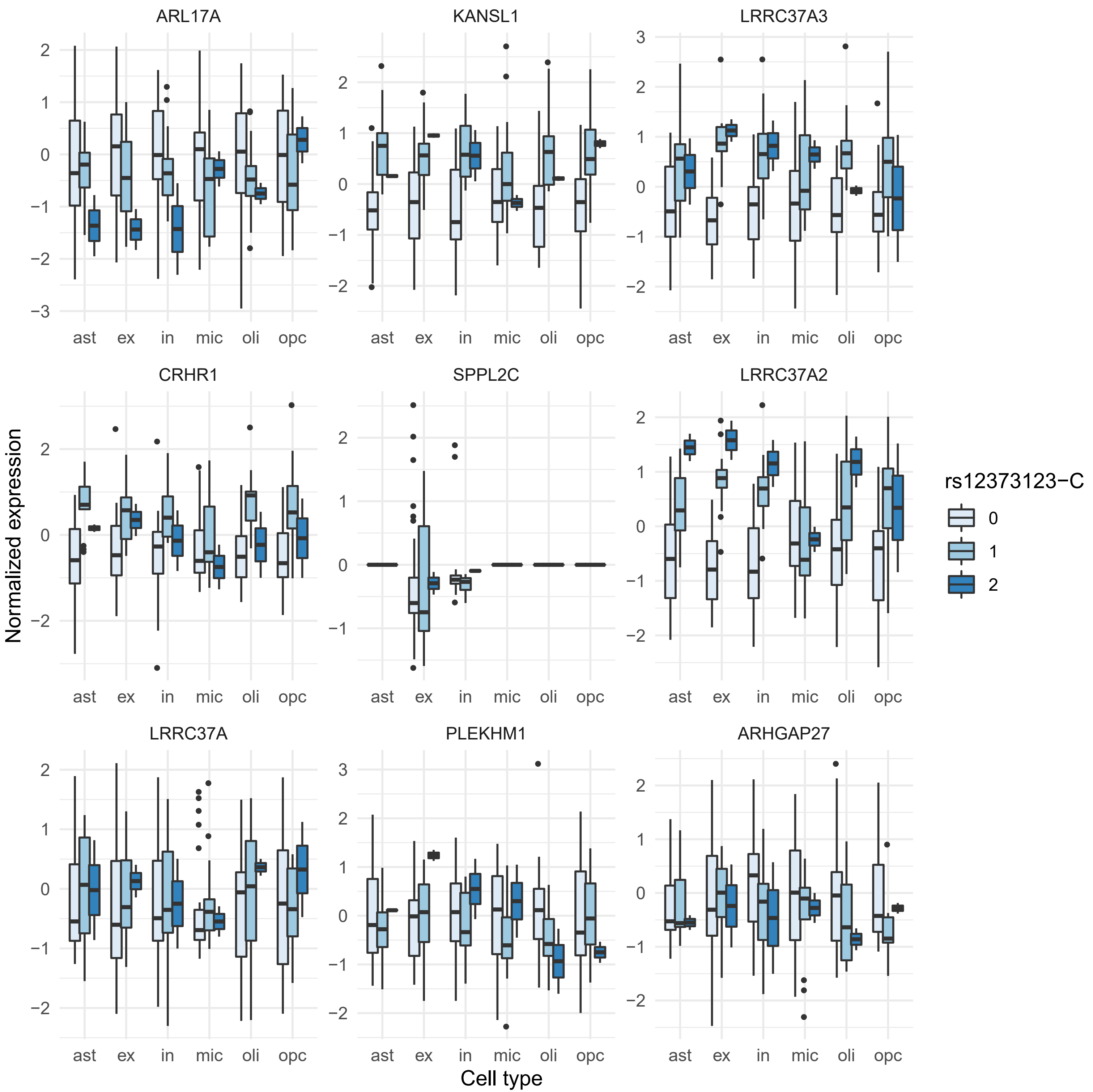

### Figure S3

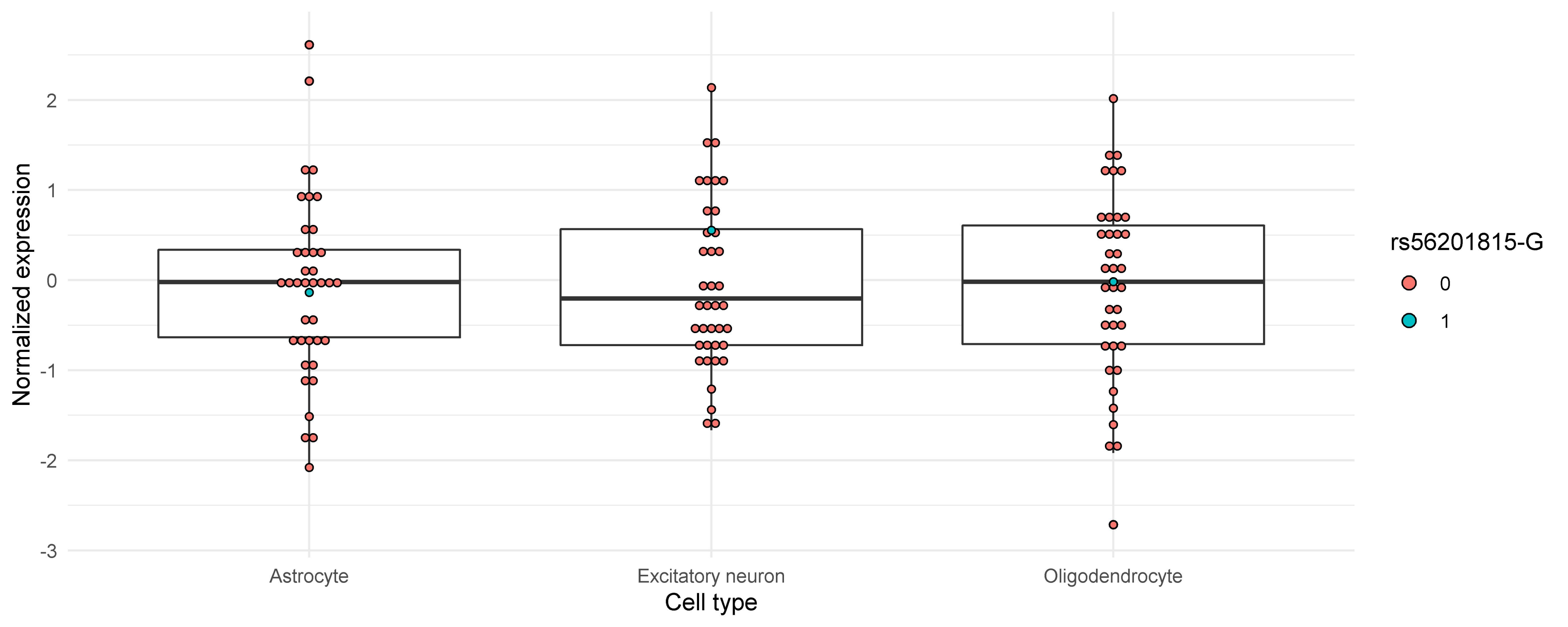

### Figure S4

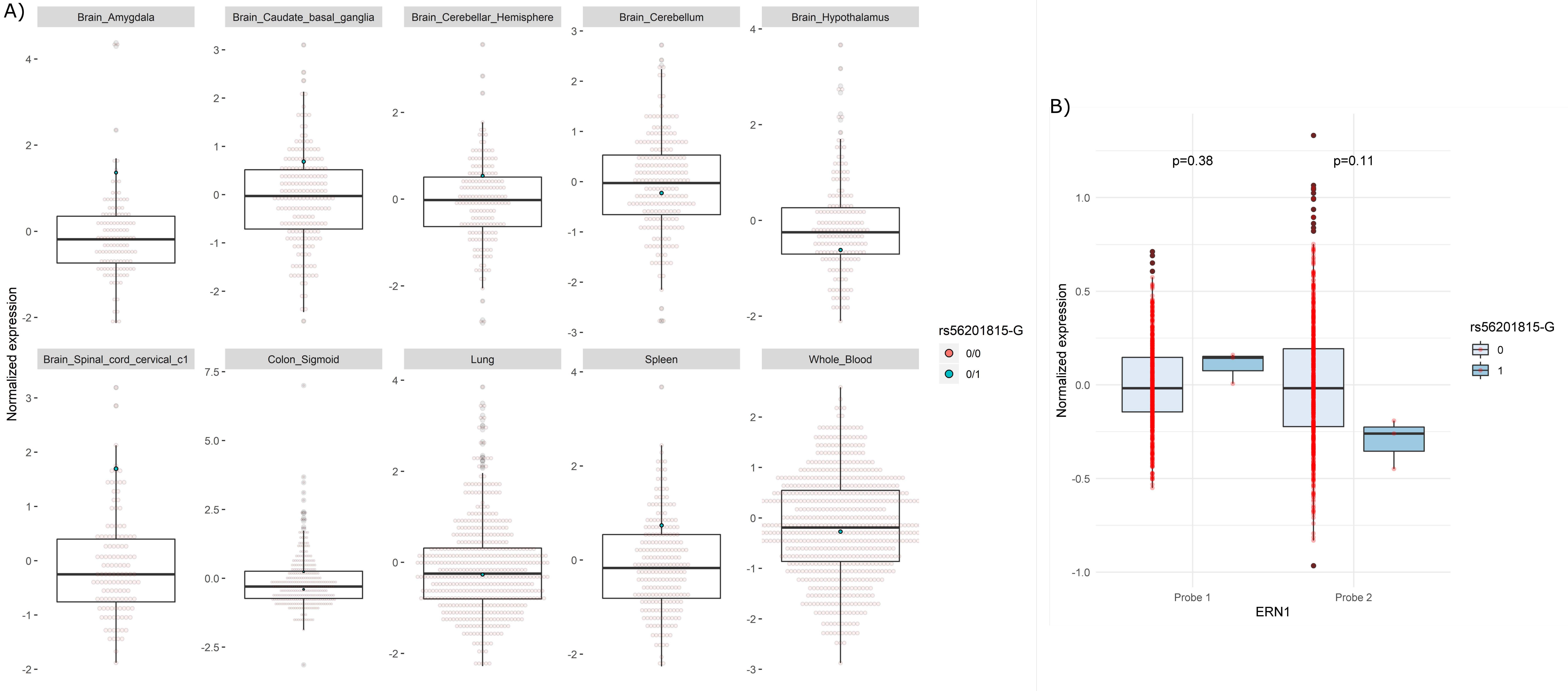

### Figure S5

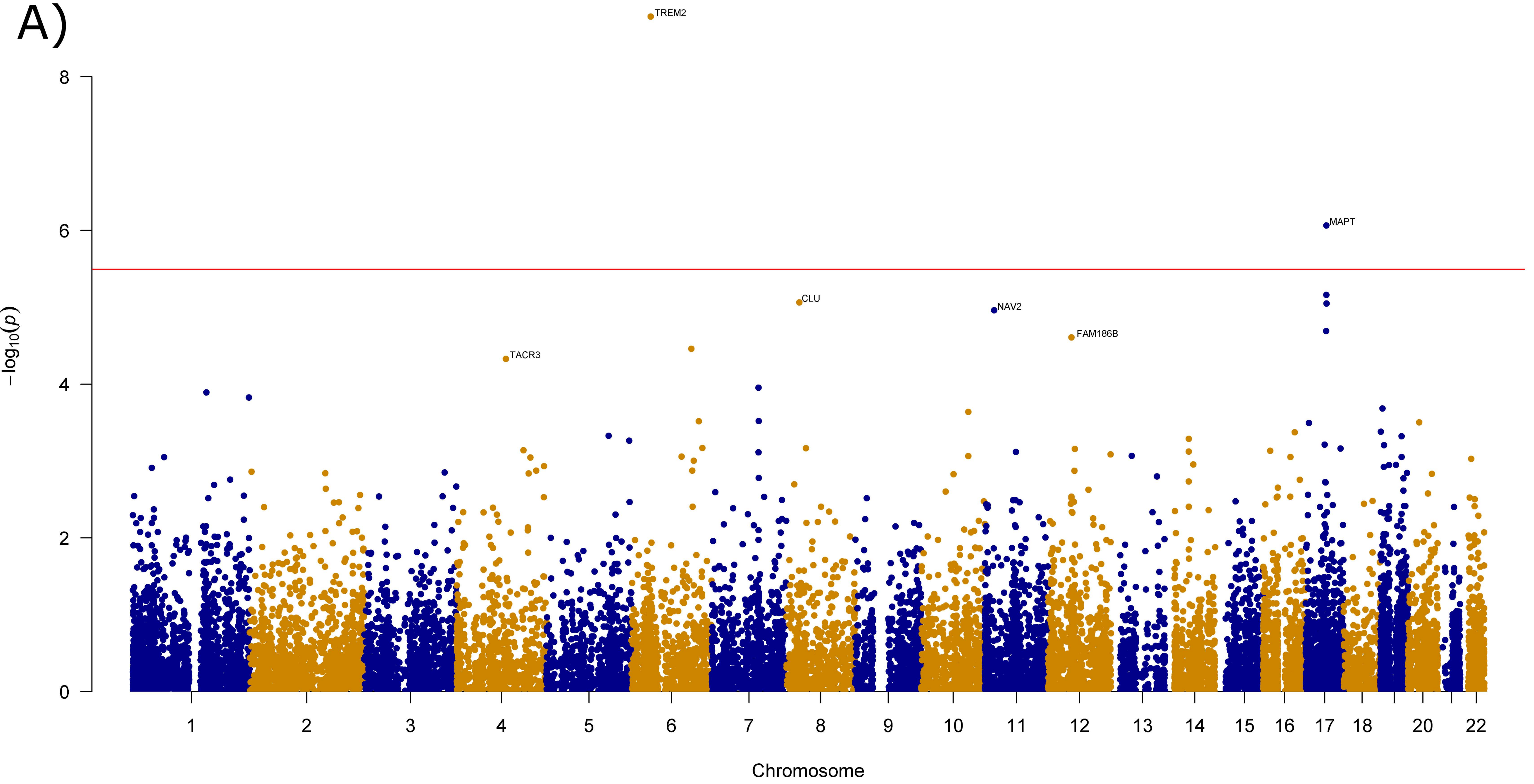

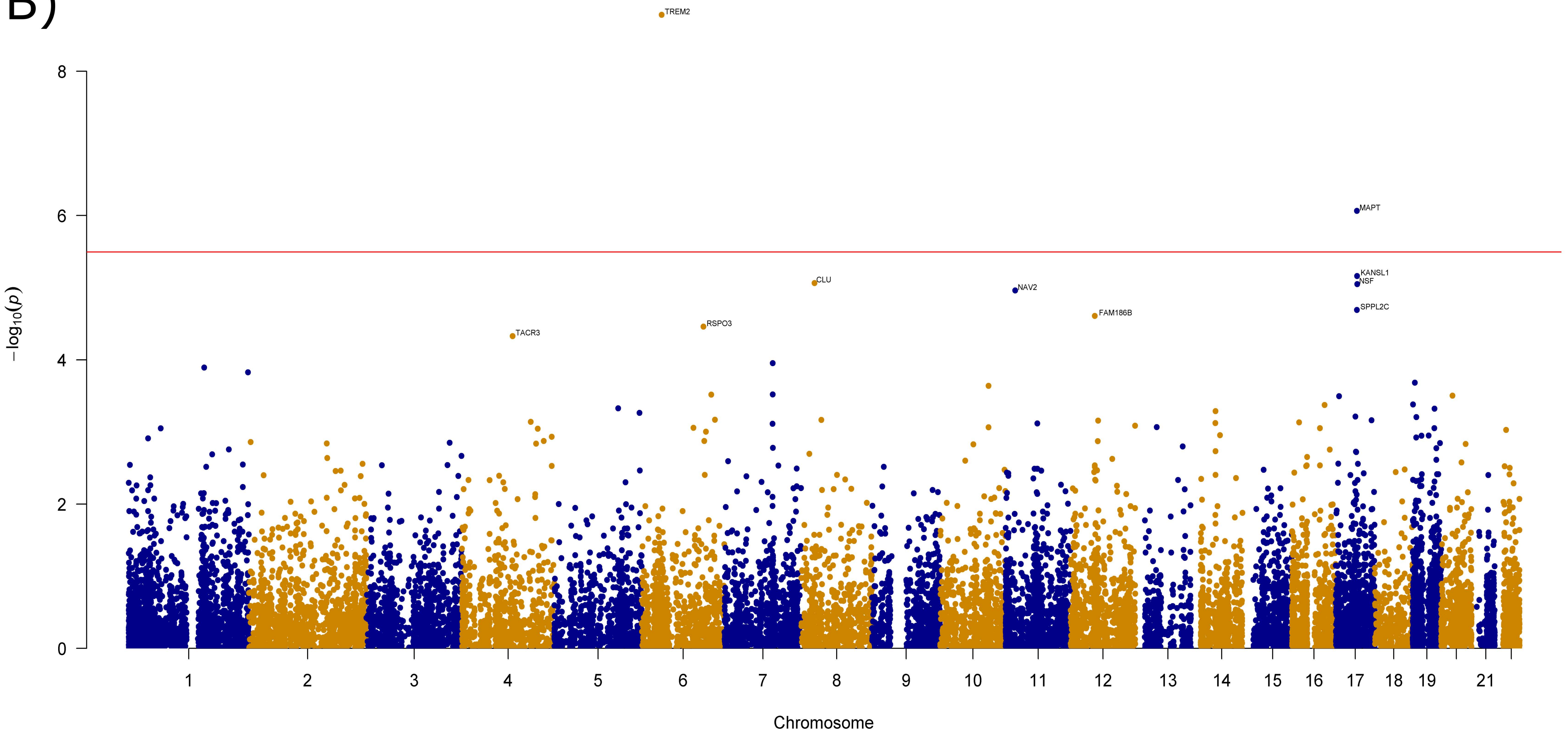
